# Experimentally disambiguating models of sensory cue integration

**DOI:** 10.1101/2020.09.01.277400

**Authors:** Peter Scarfe

## Abstract

Sensory cue integration is one of the primary areas in which a normative mathematical framework has been used to define the “optimal” way in which to make decisions based upon ambiguous sensory information and compare these predictions to behaviour. The conclusion from such studies is that sensory cues are integrated in a statistically optimal fashion. However, numerous alternative computational frameworks exist by which sensory cues could be integrated, many of which could be described as “optimal” based on different criteria. Existing studies rarely assess the evidence relative to different candidate models, resulting in an inability to conclude that sensory cues are integrated according to the experimenter’s preferred framework. The aims of the present paper are to summarise and highlight the implicit assumptions rarely acknowledged in testing models of sensory cue integration, as well as to introduce an unbiased and principled method by which to determine, for a given experimental design, the probability with which a population of observers behaving in accordance with one model of sensory integration can be distinguished from the predictions of a set of alternative models.

## Introduction

### Integrating sensory information

Humans have access to a rich array of sensory data from both within and between modalities upon which to base perceptual estimates and motor actions. These data are treated as consisting of quasi-independent sensory “cues”. Given a set of cues, the question then becomes how information is *integrated* to generate a robust percept of the world (Ernst & Bulthoff, 2004). Mathematically, there are multiple ways in which this could occur (Jones, 2016; Tassinari & Domini, 2008; Trommershauser et al., 2011), however, currently the most popular theory is the minimum variance unbiased estimator model (MVUE). MVUE forms part of broader computation frameworks such as modified weak fusion (MWF) (Landy et al., 1995) and is related to Bayesian models of sensory perception (Knill & Richards, 1996). Indeed, sensory cue integration has been described as the “… poster child for Bayesian inference in the nervous system” (Beierholm et al., 2009, p. 1).

In MVUE, given two cues, *Ŝ*_*A*_ and *Ŝ*_*B*_, each corrupted by statistically independent zero-mean Gaussian noise with variances 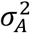 and 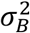, it can be shown, given some additional assumptions, that the integrated cues estimate, *Ŝ*_*C*_, is given by a simple weighted average (for derivations see Cochran, 1937; Oruc et al., 2003).

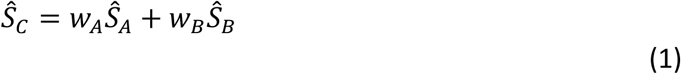

The weights are determined by the relativity reliability of the cues (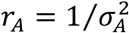 and 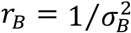) such that *w*_*A*_ = *r*_*A*_/(*r*_*A*_ + *r*_*B*_) and *w*_*B*_ = *r*_*B*_/(*r*_*A*_ + *r*_*B*_) and the standard deviation (sigma) of the Gaussian probability density function representing the integrated cues estimator is given by

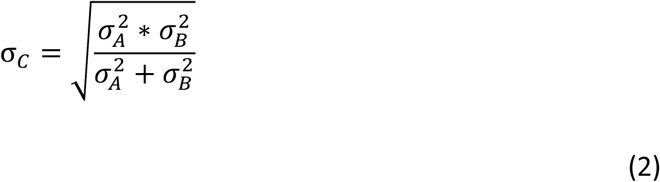

The key benefit of integrating cues in this way is that the sigma of the integrated cues estimator is always less than or equal to the sigma of the most reliable of the individual sensory cues. As a result, MVUE is often terms “optimal cue integration” (Rohde et al., 2016). Whilst there are clearly multiple benefits of combining and integrating sensory information (Ernst & Bulthoff, 2004), MVUE posits that the optimising criteria of sensory integration is to maximise the precision of the integrated cues sensory estimate. The maximum reduction in sigma (increase in precision) is achieved when the two cues are equally reliable (Figure 1). As the reliability of the cues becomes unbalanced, the increase in precision rapidly diminishes (Figure 1b). Therefore, in terms of sensitivity, for an organism to do better than simply choosing the most reliable of the two cues, the cues must be approximately matched in reliability.

**Figure 1:**
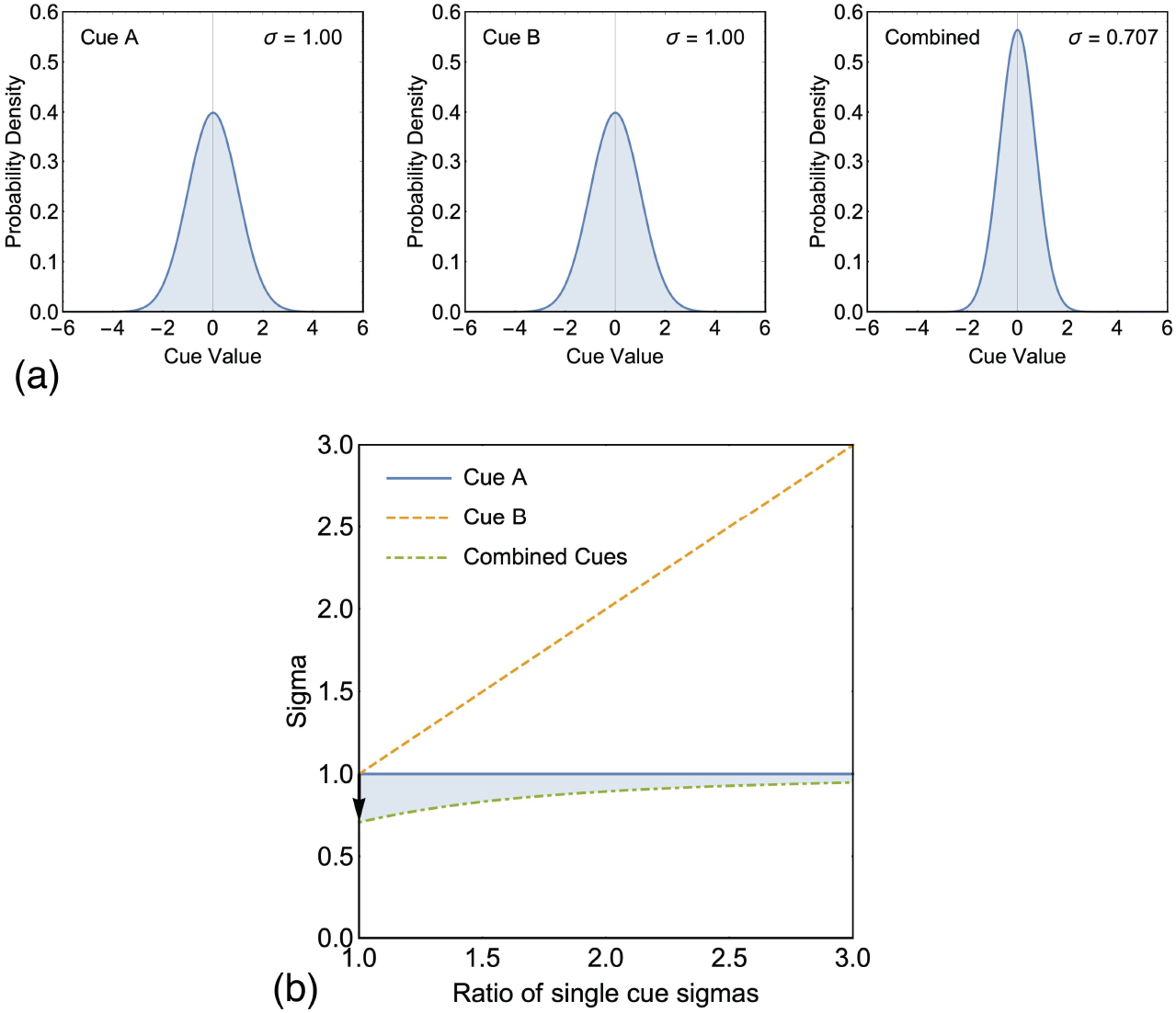
(a) Shows a hypothetical example of integrating two cues (A and B) with identical reliabilities (for both cues σ = 1). In this instance an observer would maximally benefit from combining cues in accordance with Equation 2 and obtain 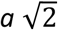 reduction in sigma. (b) Plots single cue sigmas and the integrated cues sigma associated with the two cues for a range of sigma ratios. A sigma ratio of one indicates that the two cues are equally reliable (as in (a)). A value greater than one indicates that Cue B is more variable than Cue A. The shaded region shows the increased precision afforded by integrating cues in accordance with Equation 6. The black arrow shows the maximally achievable increase in precision shown in (a).

Numerous studies purport to show that humans combine cues in accordance with MVUE (Burge, Girshick, et al., 2010; Ernst, 2006; Ernst & Banks, 2002; Gepshtein et al., 2005; Girshick & Banks, 2009; Glennerster et al., 2006; Helbig & Ernst, 2007; Hillis et al., 2002; Hillis et al., 2004; Johnston et al., 1994; Johnston et al., 1993; Knill & Saunders, 2003; Lovell et al., 2012; Saunders & Chen, 2015; Scarfe & Hibbard, 2011; Svarverud et al., 2010; Watt et al., 2005). However, for the MVUE model to apply several assumptions need to be met. Whilst acknowledged across the literature, these assumptions are rarely mentioned in experimental tests of MVUE and are often simply assumed to have been met.

### Assumptions of MVUE

To be integrated in MVUE, cues must be in common units and these modelled units need to be equivalent to the units that the observer is using to make perceptual estimates. For example, if an observer is judging surface “slant” from “disparity” and “texture” cues (Hillis et al., 2004), however information from these cues is processed in the brain, it has to result in a measures of “slant” in the same units e.g. degrees or radians. If it doesn’t, Equation 1 is meaningless as it will be averaging a property in two different units. As a result, frameworks which incorporate MVUE as a core component, such as MWF, have a “cue promotion” stage prior to averaging where cues are promoted so as to be in common units (Landy et al., 1995). Cue promotion, whilst critical, is rarely directly studied (although see Burge, Fowlkes, et al., 2010; Burge et al., 2005). Additionally, whilst it is possible to experimentally evaluate the units an observer is using to make perceptual estimates, for example, Hillis et al. (2004) examined whether their observers were estimating “slant” from disparity, or simply doing the task based upon disparity gradient, this is rarely carried out. Typically, experimenters assume from the outset that cues are in common units and that these units are equivalent to those the observers are using to make perceptual estimates.

Additionally, in MVUE cues are integrated regardless of their perceptual bias. By “bias” we mean a difference between (a) the perceptual estimate of a property of the world and (b) the actual physical value of that property. This has been termed “external accuracy” (Burge, Girshick, et al., 2010). Bias is a problem, in part, because there is no reason to assume that cues which are more reliable are also least biased. As a result, there are a mathematically definable range of circumstances where integrating cues in accordance with Equations 1 and 2 results in perceptual estimates which are more precise, but less accurate with respect to the world (Scarfe & Hibbard, 2011). As experimenters have no direct access to an observers internal perceptual estimates, cues are generally assumed to be unbiased, with any bias attributed to unmodelled cue conflicts or response bias (Watt et al., 2005).

The assumption of unbiased estimates is problematic given the large reproducible perceptual biases shown in real world environments (Bradshaw et al., 2000; Koenderink, van Doorn, Kappers, & Lappin, 2002; Koenderink, van Doorn, Kappers, & Todd, 2002; Koenderink et al., 2000; Wagner, 1985), and in expertly controlled experiments with computer generated stimuli (Watt et al., 2005). These biases suggest that sensory cues are not necessarily accurately calibrated with respect to the world (Adams et al., 2001; Henriques & Cressman, 2012; McLaughlin & Webster, 1967; Scarfe & Glennerster, 2014; Welch et al., 1993) and importantly, it has been shown that the integration of sensory cues does not lead to those same cues being accurately calibrated (Smeets et al., 2006). As a result, it is now becoming accepted that bias in sensory estimates needs to be accounted for in models of sensory cue integration (see Ernst & Di Luca, 2011). Indeed, one can experimentally examine the effects of discrepancies between cues and model cue integration in terms of causal inference, whereby the brain evaluates the probability with which signals come from a common or distinct external causes and uses this to gate cue integration (Beierholm et al., 2009; Gepshtein et al., 2005; Körding et al., 2007).

For Equations 1 and 2 to hold, each cue needs to be well represented by a statistically independent Gaussian probability density function (Cochran, 1937). A clear case where the this does not hold is for circularly distributed variables such as planar direction. With circularly distributed variables the von Mises distribution should be used, and equations similar to MVUE can be derived with some additional assumptions (Murray & Morgenstern, 2010). However, many studies simply assume that over the stimulus domain tested, Gaussian distributions provide a good enough approximation to the underlying von Mises distributions (Hillis et al., 2004). Similarly, when statistical independence does not hold, corrections to the MVUE equations can be derived to account for correlated noise (Oruc et al., 2003), however in virtually all studies, statistical independence is assumed *a priori* or the correlation assumed to be so small that MVUE provides a valid approximation.

The final assumption we consider here is that over the domain being investigated the perceptual scale of the cues is linear. Perceptual scales are known to be nonlinear (Rohde et al., 2016), so the domain over which cue integration is investigated is typically restricted and assumed to be a close approximation to linear (e.g. Hillis et al., 2004). However, even in these instances, it has been claimed that in some circumstances observers may be making perceptual estimates based upon confounding cues which are nonlinear over the experimental domain and that as a result the experimental methodology used to estimate the precision of cues will misestimate the variance of the underlying estimators (Todd et al., 2010; Todd & Thaler, 2010). This continues to be a contentious area of active debate (Saunders & Chen, 2015; Todd, 2015).

### Experimentally testing MVUE

Testing MVUE equates to seeing if the numerical predictions made by Equations 1 and 2 correspond with observer behaviour. Of the two predictions, Rohde, van Dam and Ernst (2016, p. 7) describe Equation 2 as the “essential prediction” of optimal cue integration. They point out that seeing performance line with Equation 1 is “… by itself not sufficient to show that optimal integration occurs” (Rohde et al., 2016, p. 7). As a result, “(i)f one can show only a bias in the results (equation (1)) but not a reduction in noise (equation (2)), one *cannot conclude that optimal integration occurred* …” (Rohde et al., 2016, p. 10. Note: equation numbers have been changed to correspond to the equivalent equations in the current paper and italics added.).

There are two key reasons for this. First, identical predictions to Equation 1 can be made by alternative models of perceptual processing, including those in which cues are not integrated in any way. Second, testing Equation 1 by adding an experimental cue conflict through a “perturbation analysis” (Ernst & Banks, 2002; Young et al., 1993) can be severely disrupted if one or more of the perceptual estimates is biased. To demonstrate this, following on from the example above, we can experimentally add a perturbation of value Δ to the cue *Ŝ*_*B*_, such that

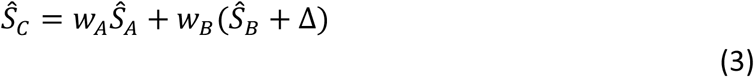

We can then ask what value of bias *β* in cue *Ŝ*_*A*_ would be required to eradicate any evidence of optimal cue integration

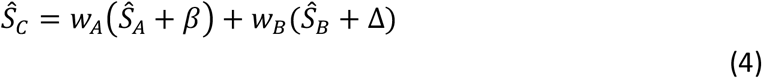

Recognising that *w*_*A*_ + *w*_*B*_ = 1 and solving for *β* gives

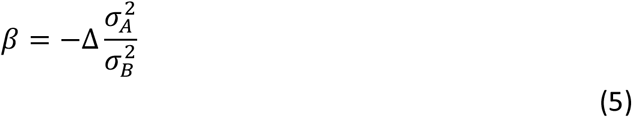

Thus, all signs of optimal cue integration can be eliminated if one or more of the perceptual estimates are biased and this depends on the relative reliability of the cues and the magnitude of the perturbation. Rohde et al. (2016) recommend that Δ be 1 to 1.5 (and no larger than 2) Just Noticeable Differences (JND), where a JND is given by 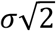, so as not to elicit cue veto (Landy et al., 1995). Therefore, whilst the ratio of cue variances being exactly that shown in Equation 5 is unlikely, cues only need to be biased by a small amount to significantly interfere determining cue weights through a perturbation analysis. This, coupled with identical predictions to Equation 1 being made by alternative models, is why Equation 2 is seen as the essential prediction of MVUE (Rohde et al., 2016).

### Comparing models of cue integration

Although MVUE is the most widely accepted model of cue integration, there are numerous alternatives, many of which take into account the reliability of sensory cues (Arnold et al., 2019; Domini & Caudek, 2009; Jones, 2016; Rosas & Wichmann, 2011; Tassinari & Domini, 2008). Much of the difference between models comes down to the computational architecture of the underlying system (Beierholm et al., 2009; Körding et al., 2007; Trommershauser et al., 2011). Therefore, as within any area of science, the question comes down to designing experiments which can distinguish between competing models. However, until recently, very few papers compared the predictions of MVUE to alternative models in any rigorous fashion (for exceptions see Acerbi et al., 2018; de Winkel et al., 2018; Lovell et al., 2012). This has been recognised as a clear weakness in claiming that cues are integrated in accordance with MVUE (Arnold et al., 2019).

An additional problem is that readers are often required to judge the fit of the data to MVUE “by eye”, without any accompanying statistics detailing the fit of the model to the data (e.g. Ernst & Banks, 2002; Hillis et al., 2004). A recent review has suggested that the adherence to MVUE, can be assessed visually and has provided a visual taxonomy of “optimal”, “sub-optimal”, “ambiguous”, “near optimal” and “supra-optimal” performance (Rohde et al., 2016, p. 23). This visual taxonomy, based on judging the fit to the predictions of MVUE from visual inspection of (1) the data, (2) the error bars around the data, and (3) the predictions of MVUE, has started to be used by researchers to assess the “optimality” of experimental data (Negen et al., 2018).

A visual taxonomy is problematic for many reasons. First, across a range disciplines, including psychology, behavioural neuroscience and medicine, leading researchers have been shown to have fundamental and severe misconceptions about how error bars relate to statistical significance and how they can be used to support statistical inferences from data (Belia et al., 2005; Cumming et al., 2007). Second, as will be seen, alternative models of cue integration provide highly correlated predictions with one another. Therefore, “eyeballing” the fit to a single model based on visual inspection is likely to lead to fundamental mistakes in inferring the extent to which a given model fits the data, especially when the literature in this area tends to be focused upon verification, but not necessarily falsification (Rosas & Wichmann, 2011). Finally, there are techniques which can be easily used to assess the fit of a set of candidate models to data in a far more objective way.

### Outline of the current study

Here we present a technique consisting of simulating end-to-end experiments (behaviour of observers in an experiment, fitting of psychometric functions, estimation of parameters from data, and final statistical analysis) which can be used to determine the probability with which a population of observers behaving in accordance with one model of sensory cue integration can distinguished from the predictions of a set of alternative models. Given its ubiquity, we focus on the extent to which the predictions of MVUE can be distinguished from two popular alternative models, (a) choose the cue with the minimum sigma (MS), and (b) probabilistic cue switching (PCS). There are numerous other models which could have been chosen for comparison (Jones, 2016), however, these two models have the benefits of (1) being conceptually similar to MVUE, (2) require experimental estimation of the same parameters, and (3) are reducible to comparably simple equations. They have also been compared to the predictions of MVUE in previous papers.

## Methods and Results

### Correlated predictions of alternative models

When choosing the cue with the minimum sigma (MS), the sigma of the integrated cues estimator is simply the sigma of the most reliable cue. Therefore, when the reliabilities of the two cues are imbalanced MS provides highly similar predictions to MVUE. This can be seen in Figure 2 where we re-plot discrimination thresholds for the visual, haptic and integrated cue estimators from Ernst and Banks (2002) (see S1). For the 0, 67 and 200% noise conditions the discrimination threshold for the integrated cues estimator is visually indistinguishable from the threshold of the most reliable of the individual cues (visual or haptic). Thus, the only condition in this paper where MS and MVUE make clearly different predictions is the 133% noise condition where the reliabilities of the two cues are nearly identical (grey rectangle).

**Figure 2:**
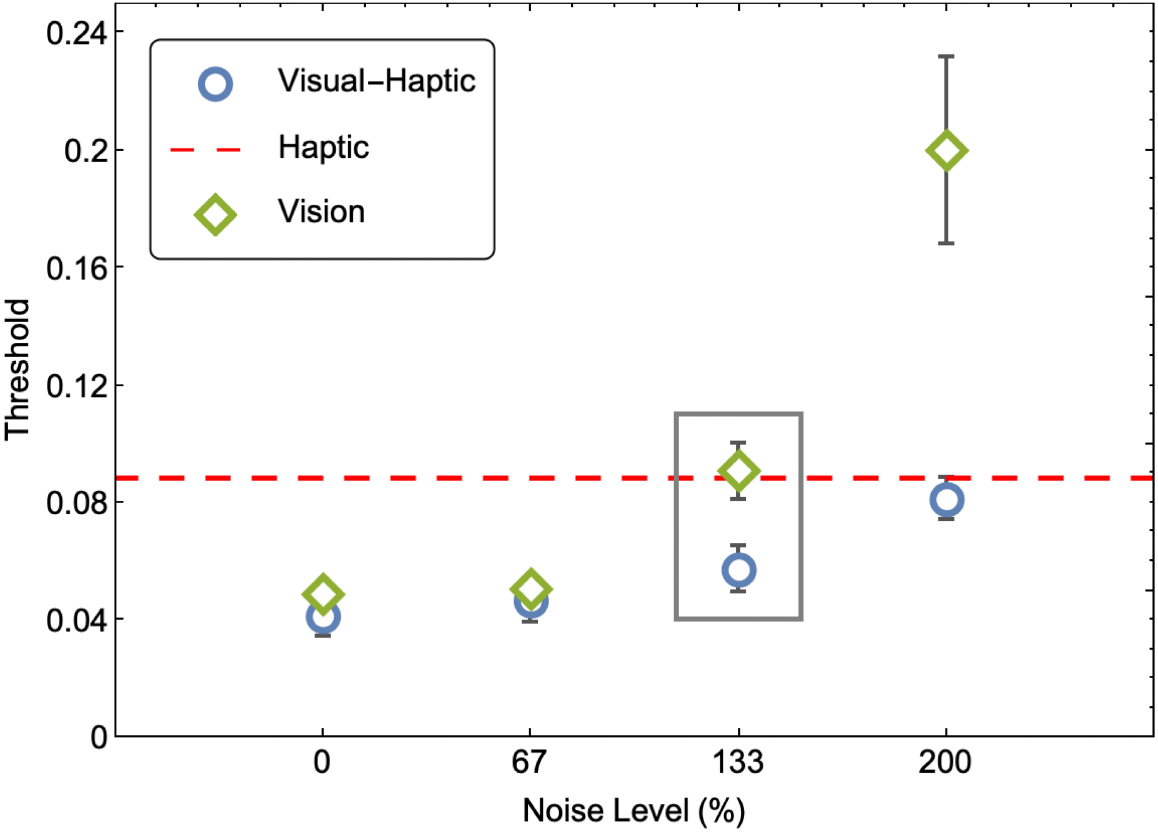
Replot of the threshold data from Ernst and Banks (2002) Figure 3d (see S1 for details). The threshold is defined as the difference between the 84% and 50% point of the underlying psychometric function. Thus, smaller thresholds represent more precise perceptual estimates. Thresholds are plotted against the % of noise in the visual modality stimulus (see Ernst & Banks, 2002 for full details). The only datapoint which can distinguish MVUE from MS is the 133% noise level stimulus (grey rectangle).

Whilst one could argue the 133% datapoints distinguish models, there are a few complications in making this inference, as there are no statistics reported to assess this difference and it is not stated what the error-bars (calculated over four observers) show. As a result, the reduction in sigma for the 133% condition is a visual judgement on the part of the reader. As detailed above, leading researchers have been shown to have fundamental misconceptions about visual judgements about statistical significance from visual inspection (Belia et al., 2005; Cumming et al., 2007). Whilst attaching a “p-value” to a result is clearly not the only way in which to make inferences from data (and can be highly problematic) (Kruschke, 2010, 2011). It is acknowledged that more rigorous methodologies are required to distinguish between competing cue integration models (Acerbi et al., 2018).

Probabilistic cue switching (PCS) (Byrne & Henriques, 2013; de Winkel et al., 2018; Nardini et al., 2008; Serwe et al., 2009) proposes that observers do not integrate cues to form a single perceptual estimate, rather, they use a single cue at a given time and switch between cues with the probabilities *p*_*A*_ and *p*_*B*_ (where *p*_*A*_ + *p*_*B*_ = 1). The mean and sigma of the integrated cues estimator is given by

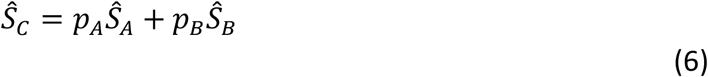

and

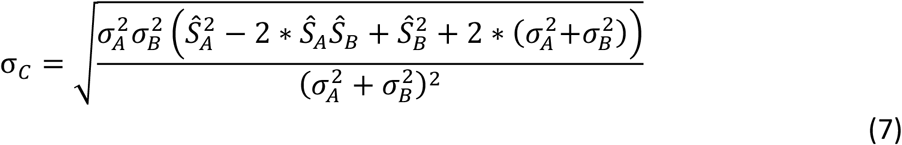

When *Ŝ*_*A*_ = *Ŝ*_*B*_, Equation 7 simplifies further to

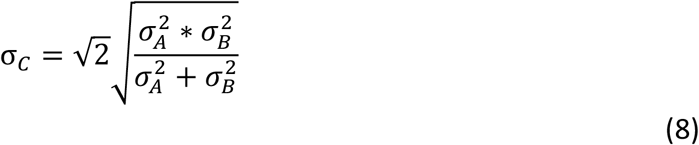

The similarities between Equations (6) and (1), and Equations (8) and (2) are clear. Note that Equations (6) and (1) provide identical predictions when *p*_*A*_ = *w*_*A*_ and *p*_*B*_ = *w*_*B*_. In other words, for the mean of the integrated cues estimator, a model in which cues are not integrated and instead used completely independently can produce identical predictions to MVUE. Throughout the paper where PCS is modelled, we have set *p*_*A*_ = *w*_*A*_ and *p*_*B*_ = *w*_*B*_, so as to be consistent with previous research where these parameters are estimated and modelled (e.g. Byrne & Henriques, 2013). However, it is true that in reality *p*_*A*_, *p*_*B*_, *w*_*A*_, and *w*_*B*_ could be determined by more than simply the relative reliability of cues as measured with a 2AFC forced choice experiment that experimenters adopt to estimate these parameters (see Jacobs, 2002 for an extended discussion).

Figure 3a-c plots the predictions for the sigma of the integrated cues estimator under MVUE, MS and PCS for a range of cue relative reliabilities. For PCS, the two cues have been set to have the same mean (i.e., Equation 8). The three models provide highly correlated predictions. Figures 3d and 3e take the difference in the predictions of the models. MS and PCS both provide maximally different predictions from MVUE when the sigma of the individual cues is identical (positive diagonal), and the absolute magnitude of this difference increases with the sigma of the two cues (compare the bottom left to top right in each plot). Also plotted are data points from two of the most widely cited papers on optimal cue integration, Ernst and Banks (2002) and Hillis, Watt, Landy and Banks (2004). Whilst some of these datapoints lay near the positive diagonal, many datapoints fall into areas of the parameter space which poorly distinguished MVUE from MS and PCS.

**Figure 3:**
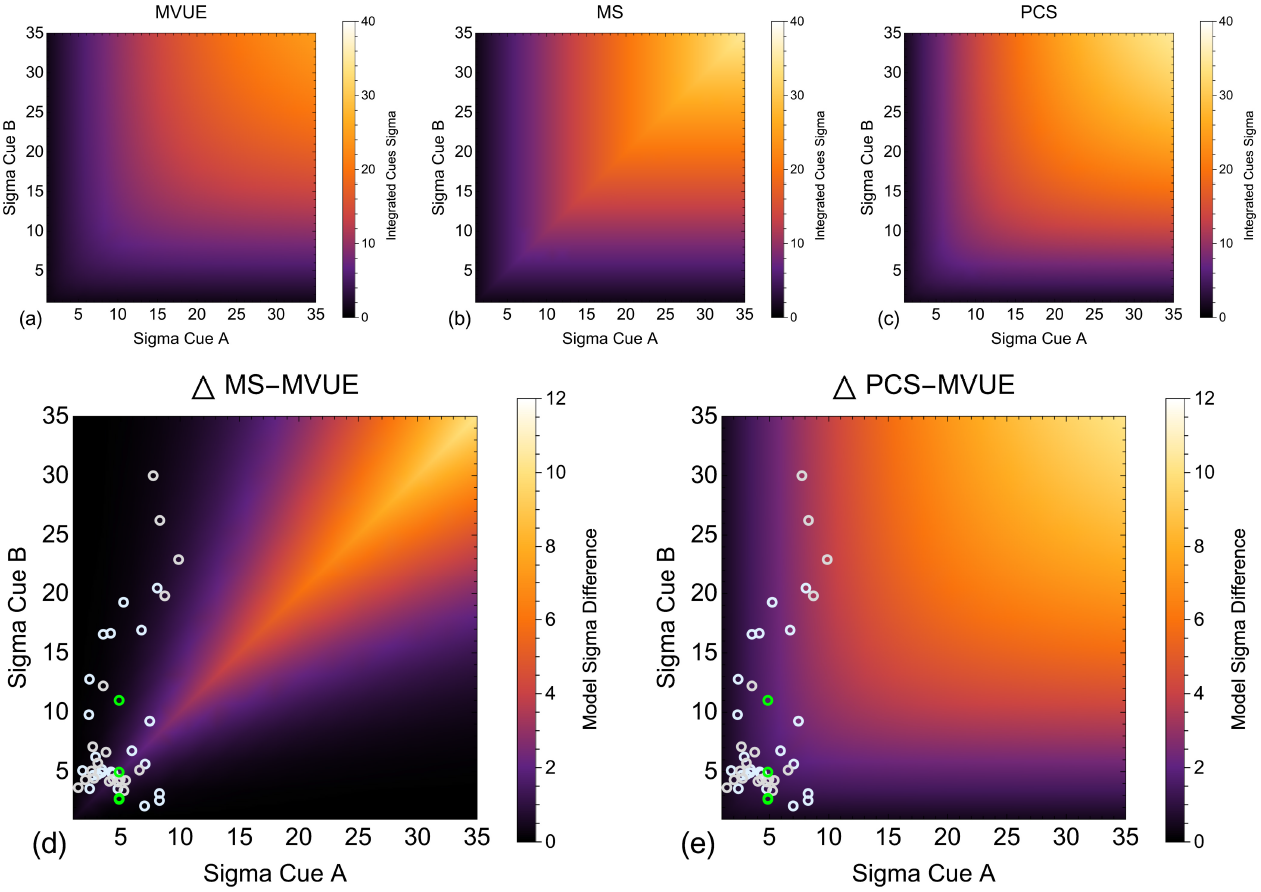
Shows the integrated cues sigma, for a range of two-cue sigma values under our three models of cue integration, (a) MVUE (Equation 2), (b) MS and (c) PCS (Equation 8). (d) Plots the difference in integrated cues sigma predicted by MS versus MVUE and (e) PCS versus MVUE. In these two plots Green symbols show the sigma values from Figure 3d in Ernst and Banks (2002) for the perception of object height. Cyan and grey symbols show sigma values from Figure 11 in Hillis, Watt, Landy and Banks (2004) for the perception of surface slant (cyan symbols observer JMH and grey symbols observer ACD).

The correlated predictions of models of models of cue integration is a known problem (Arnold et al., 2019; de Winkel et al., 2018). Indeed, the explicit aim of some of the key earlier studies in the area was to distinguish between models. For example, Ernst and Banks (2002) aimed to distinguish between MVUE and the prevailing wisdom that vision dominated haptics when the modalities were in conflict (for an extended discussion see Rohde et al., 2016). Subsequent studies have focused on examining areas of the parameter space which maximally distinguish between models such as MVUE and MS (e.g. Takahashi et al., 2009) and more rigorous model comparison approaches have been adopted (Acerbi et al., 2018; Lovell et al., 2012), however this is not the norm. As a result, if the aim is to distinguish between models, there are many things that could be improved upon in this area.

### General methods

All simulations were carried out in MATLAB (2020a) (MathWorks, Natick, MA, USA) on an 8-Core Intel Core i9 processor in a MacBook Pro running macOS 10.15. The simulations reported were computationally expensive, so where possible they were distributed over the computer’s CPU cores using MATLAB’s Parallel Processing Toolbox. The Palamedes toolbox was used to parametrically simulate observers and fit psychometric functions (Kingdom & Prins, 2010, 2016; Prins & Kingdom, 2009, 2018).

### Simulation Set 1: Effects of relative reliability and number of observers in an experiment on distinguishing between candidate models

#### Methods

##### Simulating observers

Observers were assumed to have access to two cues (*Ŝ*_*A*_ and *Ŝ*_*B*_) from which to make an integrated cues perceptual estimate (*Ŝ*_*C*_) about a property of the world. The mean of the cues prior to any perturbation was the same (55mm as in Ernst and Banks (2002)). Cue A always had the same sigma (σ_*A*_ = 4.86), which is approximately that of the haptic cue in Ernst and Banks (2002). Cue B had a sigma given by σ_*B*_ = σ_*A*_*r* where *r* varied between 1 and 4 in 27 linearly spaced steps. Whilst it has been suggested that to test for optimal cue integration the sigma ratio should be no larger than 2 (Rohde et al., 2016, p. 15), it is evident that experimenters go beyond this reliability ratio (Figure 3). Thus, in the simulations presented we go beyond a ratio of 2 to be consistent with the experimental literature. For each reliability ratio we simulated experiments where there were 4 through 30 (in steps of 1) participants. Cue integration experiments normally have few observers per experiment, but a substantial amount of data collected per observer (Rohde et al., 2016). For example, Ernst and Banks (2002) and Hillis, Watt, Landy and Banks (2004), each used four observers. Our highest observer number therefore represents an upper limit to the observers one might reasonably expect to see in a cue integration study.

The procedure described was repeated for three levels of cue conflict and four data collection regimes. The simulated conflicts, Δ, were 0, 3 and 6mm (as in Ernst and Banks (2002)). Conflicts were added by perturbing each cue by opposite amounts equal to half of the total cue conflict, that is *S*_*A*_ = 55 + Δ/2 and *S*_*B*_ = 55 − Δ/2. Estimated from the data of Ernst and Banks (2002), the above zero conflicts represented approximately 0.8 and 0.4 JNDs, which is around the recommended magnitude of cue conflict to use in a perturbation analysis (Rohde et al., 2016). In Ernst and Banks (2002) there were conditions with equal and opposite cue conflicts applied in order avoid perceptual adaptation. We did not replicate this here as our simulated observers have no mechanisms of adaptation.

We simulated performance and estimated three psychometric functions for each observer in each experiment. Two single cue functions, corresponding to the stage at which an experimenter estimates single cue sensitivities and an integrated cues condition where observers behaved in accordance with MVUE. Observers were simulated with a Cumulative Gaussian function consistent with the underlying mean and sigma of the Gaussian probability density function representing the internal estimator. Functions were sampled with the method of constant stimuli, under four data collection regimes. The method of constant stimuli was selected as this is the most widely used procedure for estimating a psychometric function. Rohde, van Dam and Ernst describe it as “… the simplest and least biased method to measure a complete psychometric function”. (p.15).

The sampling space over which the psychometric function was estimated was set to 20mm (based upon that of Ernst and Banks (2002)) and was always centred upon the true mean of the psychometric function. Centring on the true mean represents a best-case scenario for estimating the (normally unknown) function parameters. In terms of sampling density, Rohde et al. (2016) conclude that “(i)n most cases a fixed set of seven or nine comparison stimuli can be identified that suits most observers” (p. 14). Here we adopt the upper of these suggestions and spaced the stimuli linearly across the sampling range.

It is an open question how many times each stimulus should be sampled. Rohde, van Dam and Ernst (2016) suggest that when the mean and slope of the function need to be estimated around 150 trials should be used. Kingdom and Prins (2010) suggest that “400 trials is a reasonable number to aim for when one wants to estimate both the threshold and slope of the PF” (p. 57. PF, being Psychometric Function). In a simulation study, Wichmann and Hill (2001a) found that for some of their simulated sampling schemes 120 samples in total per function was often “… too small a number of trials to be able to obtain reliable estimates of thresholds and slopes …” (p. 1302). Therefore, here, in separate simulations, we examined sampling with 10, 25, 40 and 55 trials per stimulus level, giving us 90, 225, 360, and 495 trials per function.

Piloting showed that throughout the present study, these parameters resulted in well fit psychometric functions (see S2 and the criteria adopted for rejected functions detailed below). Whilst not as widely used, we could have used an adaptive method by which to sample the psychometric function (Leek, 2001). We opted not to do so (a) to be consistent with the most widely used psychophysical methods used in the literature (Rohde et al., 2016), (b) to avoid decisions related to which method to use (Kingdom & Prins, 2016; Kontsevich & Tyler, 1999; Pentland, 1980; Prins, 2013; Watson, 2017; Watson & Pelli, 1983), and (c) to avoid the issue to the adaptive method getting “stuck” in uninformative regions of the parameter space (see Prins, 2013). Additionally, adaptive procedures are used when the experimenter does not know the parameters of the underlying functions, which was not the case here.

##### Fitting functions

With the above permutations, for the first set of simulations, we simulated 27 (reliability ratios) x 27 (number of observers / experiment) x 4 (data collection regimes) x 3 (cue conflicts) x 100 (repetitions of experiment) = 874800 experiments. In total these experiments contained 14871600 simulated observers. Simulated data was fit with Cumulative Gaussian functions by maximum likelihood using the Palamedes toolbox. Whilst other fitting methods could be used, for example, fitting based on a Bayesian criterion (Kingdom & Prins, 2010; Kuss et al., 2005; Schütt et al., 2016), fitting by maximum likelihood is currently the most widely used technique in the literature (Kingdom & Prins, 2010; Wichmann & Hill, 2001a, 2001b). For all simulations we modelled observers as making zero lapses, so when fitting functions, we fixed the lapse rate to be zero (Prins, 2012; Wichmann & Hill, 2001a, 2001b).

For our simulated observers each perceptual judgement is statistically independent of all others. Therefore, there was no need here to correct for “non-stationarity” in observers’ behaviour during the fitting process (Fründ et al., 2011; Schütt et al., 2016). This is clearly not the case in an experimental setting, where there is clear evidence that the decisions made by an observer on given trial can be influenced by previous decisions the observer has made (Fischer & Whitney, 2014; Fründ et al., 2011; Kiyonaga et al., 2017; Lages & Jaworska, 2012; Liberman et al., 2014; Liberman et al., 2018; Liberman et al., 2016; Xia et al., 2016).

The mean and standard deviation of the fitted functions were taken as the experimental estimates of the observers’ true internal parameters. In cases where a function could not be fit due to the simulated data being (1) at or around chance performance across all stimulus levels, or (2) a step function, the data for that simulated observer were removed from the analysis (see also S2). Overall, this represented 0.047% of the data. The removed observers for each number of “trials per psychometric function” were: 90 trials / function = 0.183%, 225 trials / function = 0.0025%, 360 trials / function = 0.00006%, and 495 trials / function = 0%. An alternative analysis where poorly fit functions are *replaced* by a newly simulated observer results in identical conclusions being made throughout the paper.

##### Comparing the data to alternative models

For each simulated observer, the mean and sigma of the single cue function with the lowest sigma was taken as the experimental prediction for MS. For PCS, the mean and sigma of the single cue functions were entered into Equations 6 and 7 to provide predictions for the integrated cues function, with *p*_*A*_ = *w*_*A*_ and *p*_*B*_ = *w*_*B*_. Dividing sigma’s by 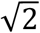, as in a typical two interval forced choice procedure (Green & Swets, 1974), was not needed as the functions were parametrically simulated. For each simulated experiment, the MVUE data were entered into a one-sample within-subjects t-test and compared to the point predictions of MS and PCS. The mean value of the alternative model prediction across observers was taken as the point prediction for each model.

Our simulated data are measured on a ratio scale and all observations are independent of one another, however we do not know that the data are normally distributed and that parametric statistical tests are appropriate. Examining the literature, it is clear that where statistical tests are run, data normality is typically not reported, but parametric statistical tests are used. Indeed, given the small number of observers in a cue integration experiment, it would be difficult to reliably estimate the normality of the data. Adopting parametric tests was therefore considered a reasonable choice (using a non-parametric Wilcoxon signed rank test results in the same conclusions being made throughout). We adopted the standard (but arbitrary) p<0.05 level for “statistical significance”.

##### Group analysis: integrated cues sensitivity

First, we examine the extent to which MVUE, MS and PCS can be distinguished based on the sensitivity of the integrated cues estimator. In Figure 4 the shading of each pixel represents the percentage of simulated experiments in which the results of a population of observers behaving in accordance with MVUE could be statistically distinguished from the numerical predictions of MS and PCS. Consistent with the correlated predictions of candidate models (Figure 3) as the sigma of the individual cues becomes unbalanced it becomes progressively more difficult to experimentally distinguish between MVUE and MS. This is especially apparent with the low number of observers that characterise typical cue integration experiments. As would be expected, when more data are collected per function models can be more easily distinguished. MVUE observers can be easily distinguished from the sigma value predicted by PCS across all sigma ratios and data collection regimes, as would be expected from Equations 2 and 8

**Figure 4:**
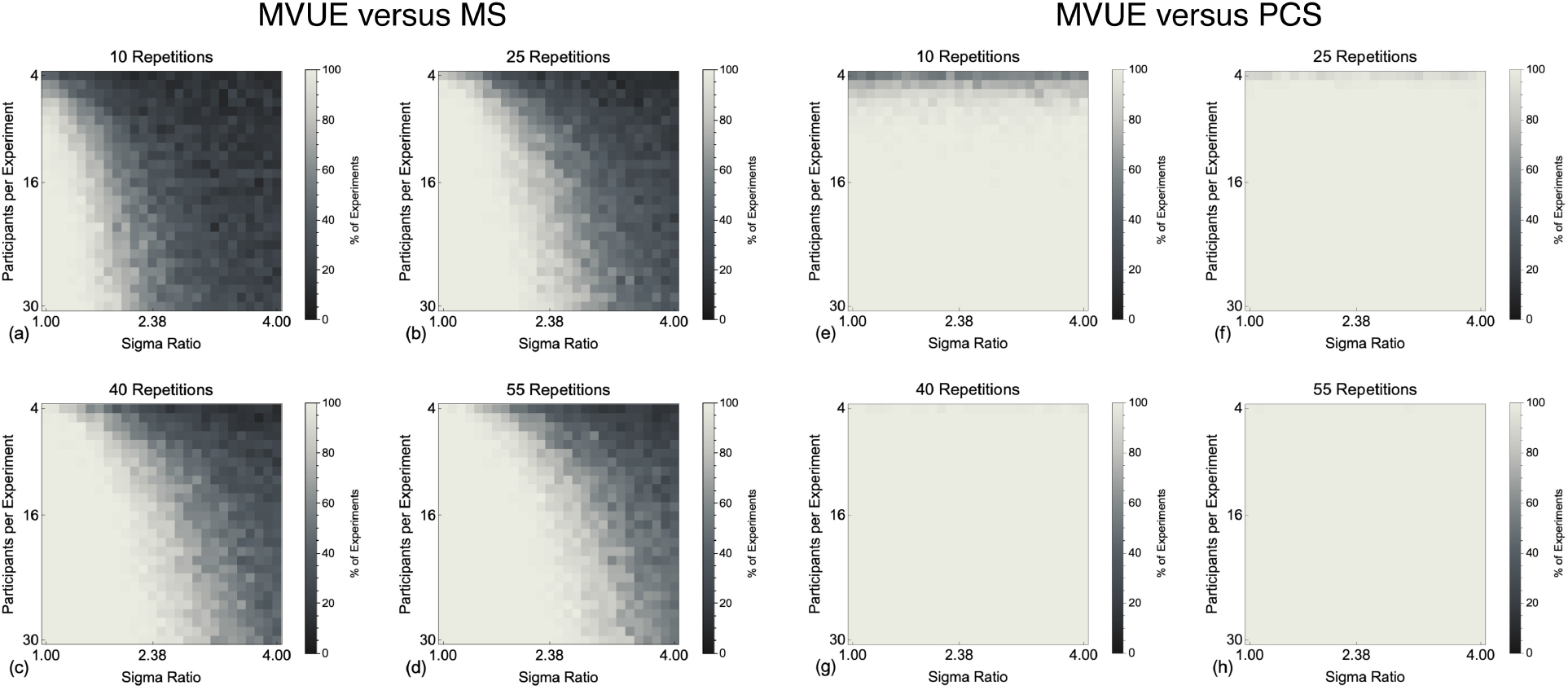
Shows the percentage of experiments in which the sigmas of the Cumulative Gaussian functions fit to our simulated population of MVUE observers could be statistically distinguished from the experimentally derived prediction of MS (a-d) and PCS (e-h). Pixels in the images show this percentage as calculated across 100 simulated experiments, for a given sigma ratio and number of participants. This is shown for (a and e) 10, (b and f) 25, (c and g) 40 and (d and h) 55, simulated trials per stimulus level on the psychometric function.

Many of the experiments shown in Figure 4 contain an unrealistically high number of observers per experiment. Therefore, Figure 5 plots the results for both comparisons, for the simulated experiments with four observers (as in Ernst and Banks (2002) and Hillis et al. (2004)). The vertical grey line shows the maximum recommended sigma ratio to use in cue integration experiments (Rohde et al., 2016), whereas the dashed grey line shows the point at which there is a 50% chance of distinguishing models. It is clear that with a representative number of observers in a typical cue integration experiment, to have any reasonable chance of distinguishing MVUE and MS, one needs to collect a large amount of data per participant and very closely match cue reliabilities. Collecting 150 trials per function across four observers with a sigma ratio of 2 would result in an approximately 25% chance of distinguishing these models, suggesting that existing guidelines (Rohde et al., 2016) may need to be improved upon. In contrast, even with four observers, PCS can be well distinguished from MVUE, for all but the lowest data collection regime.

**Figure 5:**
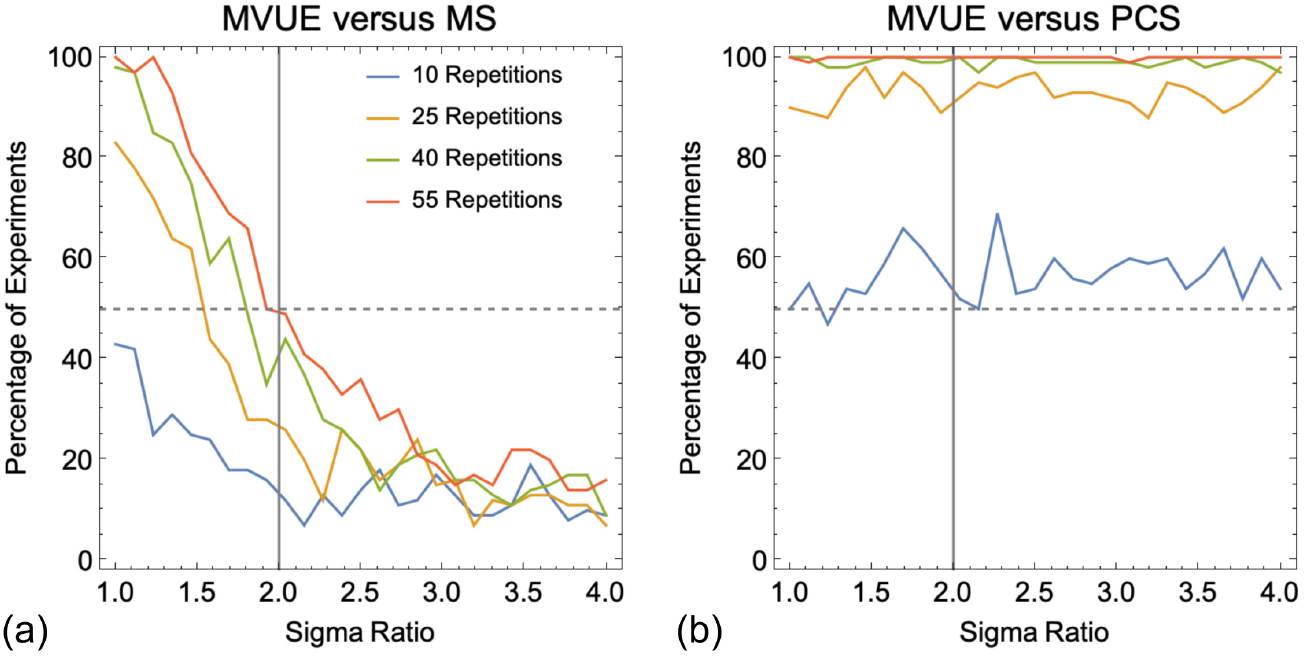
Plots the percentage of experiments in which the sigmas of the Cumulative Gaussian functions fit to a simulated population of four MVUE observers could be statistically distinguished from the experimentally derived prediction of (a) MS and (b) PCS. The dashed grey line represents the point at which there is a 50% chance of distinguishing the data from the predictions. The vertical grey line shows the maximum recommended sigma ratio to use in cue integration experiments (Rohde et al., 2016).

##### Group analysis: integrated cues percept

Next we examined the extent to which MVUE can be distinguished from MS and PCS based upon the predicted integrated cues percept when a discrepancy is experimentally introduced between cues (Young et al., 1993). With zero cue conflict the only differences in *Ŝ*_*A*_, *Ŝ*_*B*_ and *Ŝ*_*C*_ will be due to chance, so any statistical differences will represent “false positives” (see Figures S3 and S4). The false positive rate was ∼16% for MS (for 10, 25, 40 and 55 repetitions per function, the percentages are 15.99%, 16.09%, 16.21%, and 15.83%) and ∼14% for PCS (for 10, 25, 40 and 55 repetitions, the percentages are (13.62%, 13.69%, 13.82%, and 13.57%). The difference between the false positives for MS and PCS is due to the effect that the sigma of the simulated function has on the inferred mean of the function across participants. Whilst the mean and sigma of a Cumulative Gaussian functions are mathematically independent, our ability to infer these parameters by fitting psychometric functions to data is not.

Figure 6 show the data for the 3mm and 6mm cue conflicts when comparing to the predictions of MS. As with distinguishing models based on the sigmas, the ability to distinguish between models is strongly affected by the relative reliability of the cues and the data collection regime. As would be expected, the probability of distinguishing between models is greater with a larger cue conflict. Due to PCS and MVUE providing identical predictions regardless of the experimental cue conflict, the only times a population of MVUE observers are distinguishable from the predictions of PCS again represent false positives (Figures S5 and S6).

**Figure 6:**
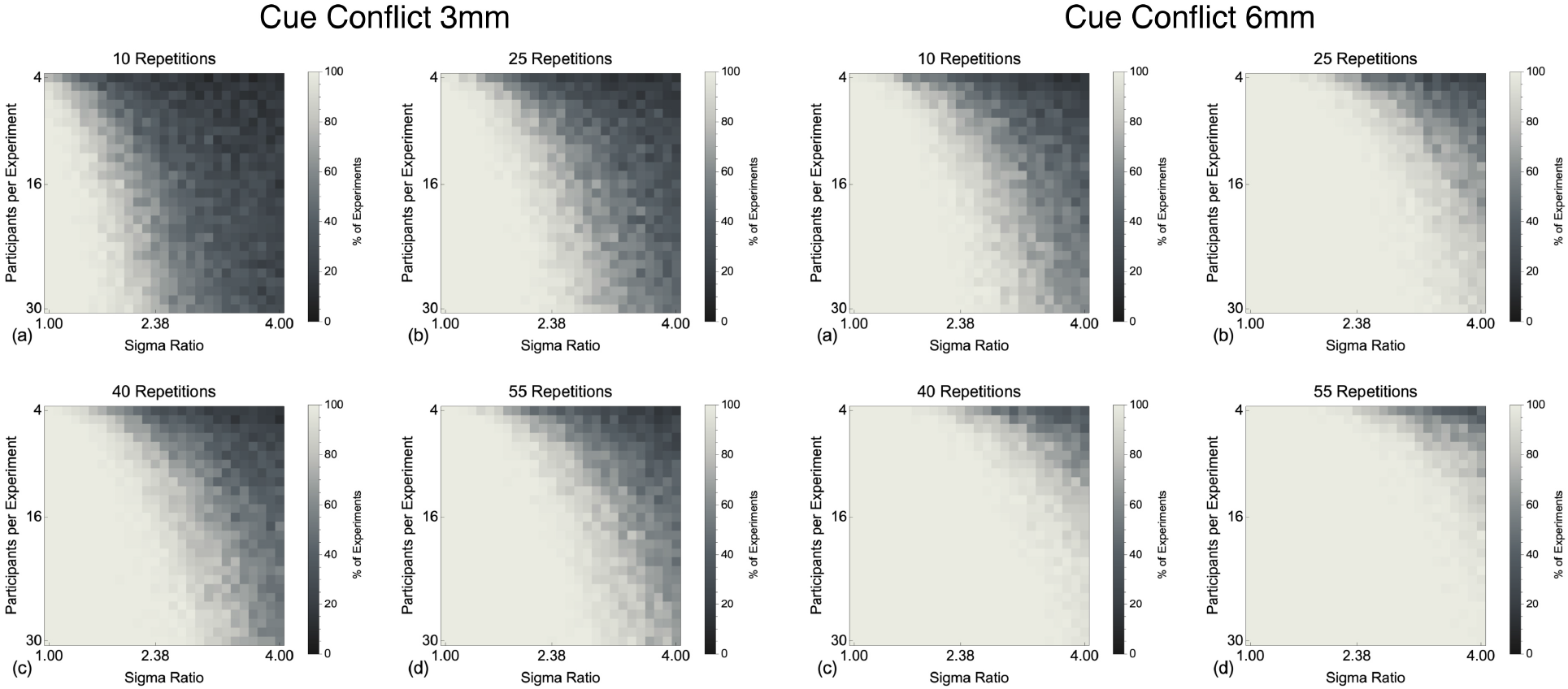
Shows the percentage of experiments in which the mean of the Cumulative Gaussian functions fit to our simulated population of MVUE observers could be statistically distinguished from the experimentally derived prediction of MS with experimental cue conflicts of 3mm (a-d) and 6mm (e-h). This is shown for (a and e) 10, (b and f) 25, (c and g) 40 and (d and h) 55, simulated trials per stimulus level on the psychometric function.

In Figure 7 we show the ability to experimentally distinguish between MVUE and MS based upon the integrated cues percept for just the simulated experiments with four observers (Ernst & Banks, 2002; Hillis et al., 2004). With no cue conflict (Delta of 0) the false positive rate is ∼12% across all data collection regimes and sigma ratios. For both cue conflicts (Delta of 3 and 6mm), the closer the reliability of cues is matched, and the more data collected, the better one can discriminate our population of MVUE observers from the predictions of MS. For a Delta of 3mm (Figure 7b), the ability to distinguish models rapidly drops off within the range of sigma ratios acceptable for a cue integration experiment (Rohde et al., 2016), such that with a sigma ratio of 3 and above, performance is comparable to that of the false positive rate (Figure 7a). By comparison, with a Delta of 6mm, within the range of sigma ratios acceptable for a cue integration experiment the ability to discriminate between models is good, with performance dropping substantially for only the most minimal data collection regime.

**Figure 7:**
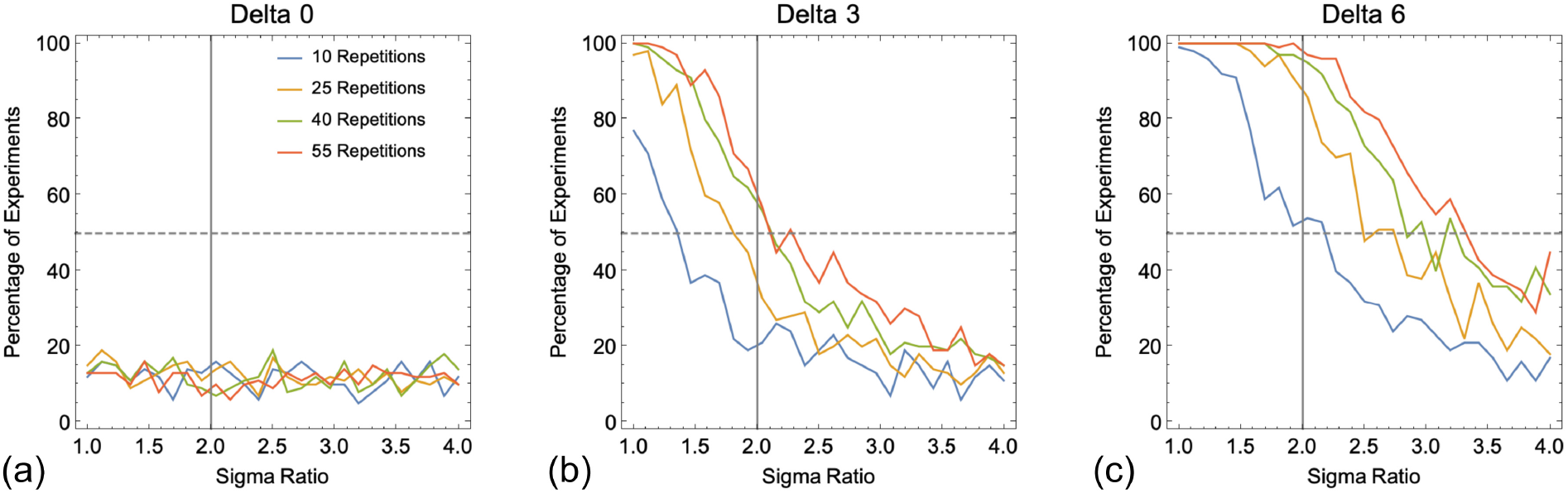
Plots the percentage of experiments in which the PSE’s of the Cumulative Gaussian functions fit to a simulated population of four MVUE observers could be statistically distinguished from the experimentally derived prediction of MS. The dashed grey line represents the point at which there is a 50% chance of distinguishing the data from MS. The vertical grey line shows the maximum recommended sigma ratio to use in cue integration experiments (Rohde et al., 2016).

One of the most striking things about the analysis presented is just how rapid the drop-off in an experimenter’s ability to distinguish a population of MVUE observers from the predictions of MS is, as the reliability of cues becomes unmatched. MVUE observers are easily distinguished from PCS in terms of the cue reliability, but impossible to distinguish based upon the integrated cues percept. MVUE observers can be more easily distinguished from MS based upon the integrated cues percept, but only dramatically so for larger cue conflicts. However, distinguishing models based upon the integrated cues percept alone is *not* sufficient to demonstrate that observers are behaving in accordance with MVUE (Rohde et al., 2016).

### Simulation Set 2: Using variation across experimental observers to distinguish between models

In the second set of simulations, we examined the case where individual observers in an experiment had different relative cue reliabilities. This is a weaker form of testing MVUE as data collection can occur in regions of the parameter space which poorly distinguishes between models (Figure 3), but it is more representative of a typical cue integration experiment where there may be variation in cue reliabilities across observers (Hillis et al., 2004; Scarfe & Hibbard, 2011) and properties of the stimuli may naturally (Hillis et al., 2004) or artificially (Ernst & Banks, 2002; Helbig & Ernst, 2007) be used to modulate the relative reliability of cues.

#### Methods

For these simulations we focused on comparing MVUE and MS as these models can be distinguished based upon both the integrated cues percept and its precision. Observers were simulated as having access from two cues (*Ŝ*_*A*_ and *Ŝ*_*B*_) from which to make an integrated cues perceptual estimate (*Ŝ*_*C*_). These cues were in conflict such that *S*_*A*_ = 55 + Δ/2 and *Ŝ* = 55 − Δ/2 (in separate experiments, Δ was either 3 or 6mm). *Ŝ*_*A*_ always had the same sigma σ_*A*_ = 4.86, which is approximately that of the haptic cue in Ernst and Banks (2002), whereas *Ŝ*_*B*_ had a randomly determined sigma of σ_*B*_ = σ_*A*_*r* where, consistent with the recommendations of Rhode et al. (2016), *r* ∈ [0.5, 2]. To select values with equal probability between these limits, for each observer we generated a random number x_*i*_ ∈ [−1, 1], and set 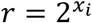. Separate simulations were run with 4, 12 and 36 observers per simulated experiment, and for 10, 25, 40 and 55 trials per stimulus level. For each combination of (a) data collection regime, (b) number of observers per experiment, and (c) cue conflict (4 × 3 × 2), we simulated 1000 experiments i.e., 32000 experiments with 416000 observers in total.

With a heterogenous population of observers the relationship between predicted and observed data are often compared using a linear regression analysis. For example, Burge, Girshick and Banks (2010) examined the perception of slant from disparity and haptic cues and reported an R^2^ of 0.60 (significance not stated) for predicted versus observed integrated cues sensitivity. Knill and Saunders (2003) also examined the perception of slant, but from disparity and texture cues, and reported R^2^ values between around 0.15 and 0.46 (p<0.05) for the predicted and observed cue weighting for different base slants. Svarverud et al. (2010) examined “texture-based” and “physical-based” cues to distance and reported R^2^ values of about 0.95 (p<0.001) for predicted and observed cue weights. The median R^2^ value in these studies is 0.53 and in all instances the authors concluded that observers were combining cues optimally in accordance with MVUE. Following these studies, a regression analysis was adopted here.

For each experiment the data from the population of observers behaving in accordance with either MVUE or MS were plotted against the predictions of each of the two candidate models. Data were fit with a first order polynomial by least squares and an R^2^ value for the fit of each model to the data calculated. Thus, there were four possible regression comparisons: (1) “MVUE v MVUE” – predictions of MVUE, plotted against data from a population of observers behaving in accordance with MVUE; (2) “MS v MS” – predictions of MS, plotted against the behaviour of a population of observers behaving in accordance MS; (3) “MVUE v MS” – predictions of the MVUE model, plotted against the data of a population of observers behaving in accordance with MS; and (4) “MS v MVUE” – predictions of the MS model, plotted against the data of a population of observers behaving in accordance with MVUE. We will refer to (1) and (2) as “consistent” predicted and observed data as the simulated data and predictions are from the same model, and (3) and (4) as “inconsistent” predicted and observed data as the simulated data and predictions arise from different models.

A set of example data (PSE and sigma) from 36 observers behaving in accordance with MVUE (with 55 samples per stimulus value and a delta of 3mm) is shown in Figure 8a-d for the “MVUE v MVUE” and “MS v MVUE” comparisons. This example represents the upper limit of observers and data collection in a typical cue combination experiment (Kingdom & Prins, 2016; Rohde et al., 2016). Figure 8a and b plot the PSE data from the MVUE observers against the experimentally derived predictions of the two candidate models, with the green and red dashed lines show the true underlying PSE for each cue. Figure 8c and d plot the observed sigma data from the MVUE observers against the experimentally derived predictions of the two candidate models, here, the dashed red line shows the fixed sigma of Cue A and the green dashed line the minimum possible sigma for Cue B.

**Figure 8:**
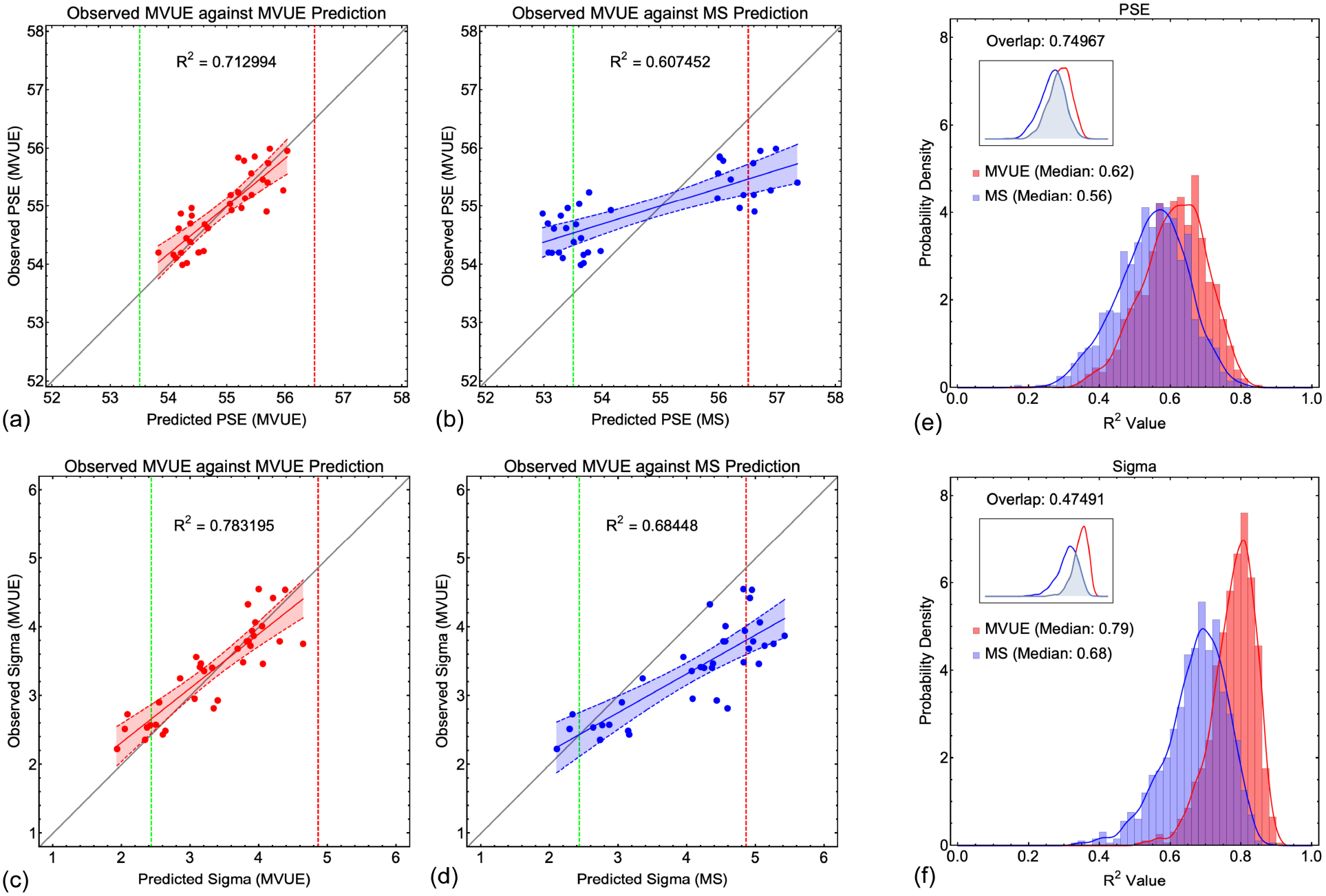
(a) through (d) show an example linear regression analysis where the data from 36 observers behaving in accordance with MVUE (with 55 samples per stimulus value and a delta of 3mm) are plotted against the predictions of the two candidate models (MVUE (a and c) and MS (b and d)), for both PSE (a and b) and Sigma (c and d). The least squares first order polynomial is shown as the solid line, with the dashed lines and shaded region showing the 95% confidence bounds around the fit. In (a) and (b) the dashed red line shows the true underlying PSE for Cue A, and the green dashed line shows the true underlying PSE for Cue B. In (c) and (d) the red dashed line shows the (fixed) sigma for Cue A, and the dashed green line the minimum possible sigma for Cue B (which varied across simulated observers). (e) and (f) show the full distributions for the R^2^ value across all 1000 simulated experiments for (e) PSE and (f) sigma. Data are shown as bar histograms and as smoothed histograms (smoothed Gaussian kernel distribution; Equation 9). Red data are from MVUE observers plotted against the predictions of MVUE; blue data are from MVUE observers plotted against the predictions of MS. The median for each data set is shown in the graphs. The inset graph shows the overlap of the smoothed histograms (Equation 10). Note that the axes of the inset graphs is smaller to ensure clarity of the overlapping region.

What is most striking from this example is that the observed R^2^ values for both PSE’s and sigmas are directly comparable to those found in the literature (and even better) *regardless* of whether the data from a population of MVUE observers were fitted with a regression against the predictions of *either* MVUE or MS. Figure 8e and f shows histograms of the observed R^2^ values for the same example, but across all 1000 simulated experiments. The raw histograms are shown overlaid with smooth kernel distributions, given by

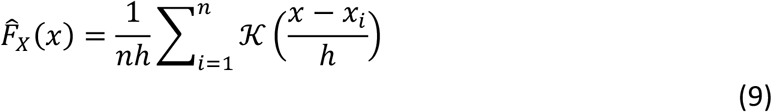

Here 𝒦 is a Gaussian kernel function, x_*i*_ ∈ [0, 1] (i.e., the domain of the R^2^ value is 0 to 1), and 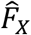 is the estimate of the unknown probability density function *F*_*x*_. The key parameter of interest is the extent to which these distributions overlap, as this determines the extent to which an R^2^ value from fitting predicted to observer data can be used to distinguish between candidate models of cue integration. The overlap of two smooth kernel distributions 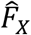 and 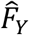 can be estimated via numerical integration (Pastore & Calcagni, 2019)

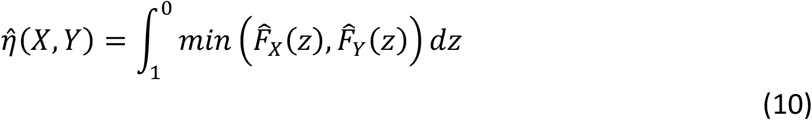

Numerically the overlap value lays between 0 (no overlap) and 1 (full overlap). This is shown inset in Figure 8e and f. As can be seen there is substantial overlap in the distribution of R^2^ values, especially so for the predicted and observed PSEs.

Data across all comparisons for both PSE and sigma are shown in Figures 9, 10 and 11. As one would expect, with more data collected per function and more observers per experiment the R^2^ values improve, with a maximal median of ∼0.7-0.8. Problematically, this pattern is present *regardless* of whether one is plotting consistent predicted and observed data (MVUE v MVUE and MS v MS), or inconsistent predicted and observed data (MVUE v MS and MS v MVUE). Across all plots there is the large overlap in the distributions of R^2^ values when plotting “consistent” and “inconsistent” predicted and observed data. With fewer observers per experiment (4 and 12 versus 36) the overlap increases greatly, to the extent that with four observers per experiment the data have near complete overlap.

**Figure 9:**
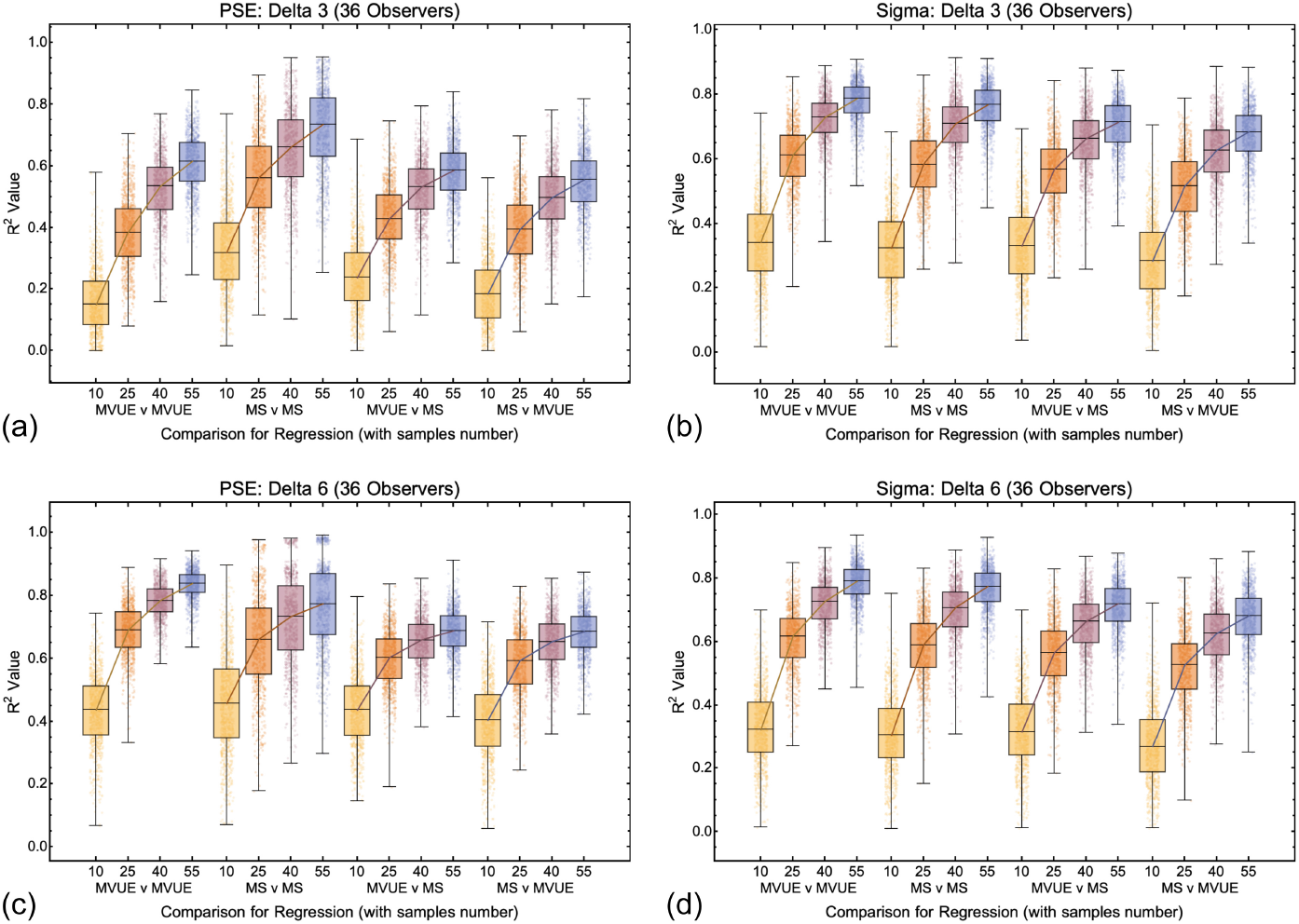
Box and whisker plots showing the distribution of R^2^ values for all conditions and comparisons in which there were 36 simulated observers per experiment for the 3mm (a and b) and 6mm (c and d) cue conflict (delta) conditions. The central box line shows the median (also shown as a line connecting the boxes), the limits of the boxes show the 25% and 75% quantiles and the limits of the bars (whiskers) show the maximum and minimum values. Also shown are all 1000 datapoints per condition (dots). For increased clarity the dots have been randomly jittered laterally.

**Figure 10:**
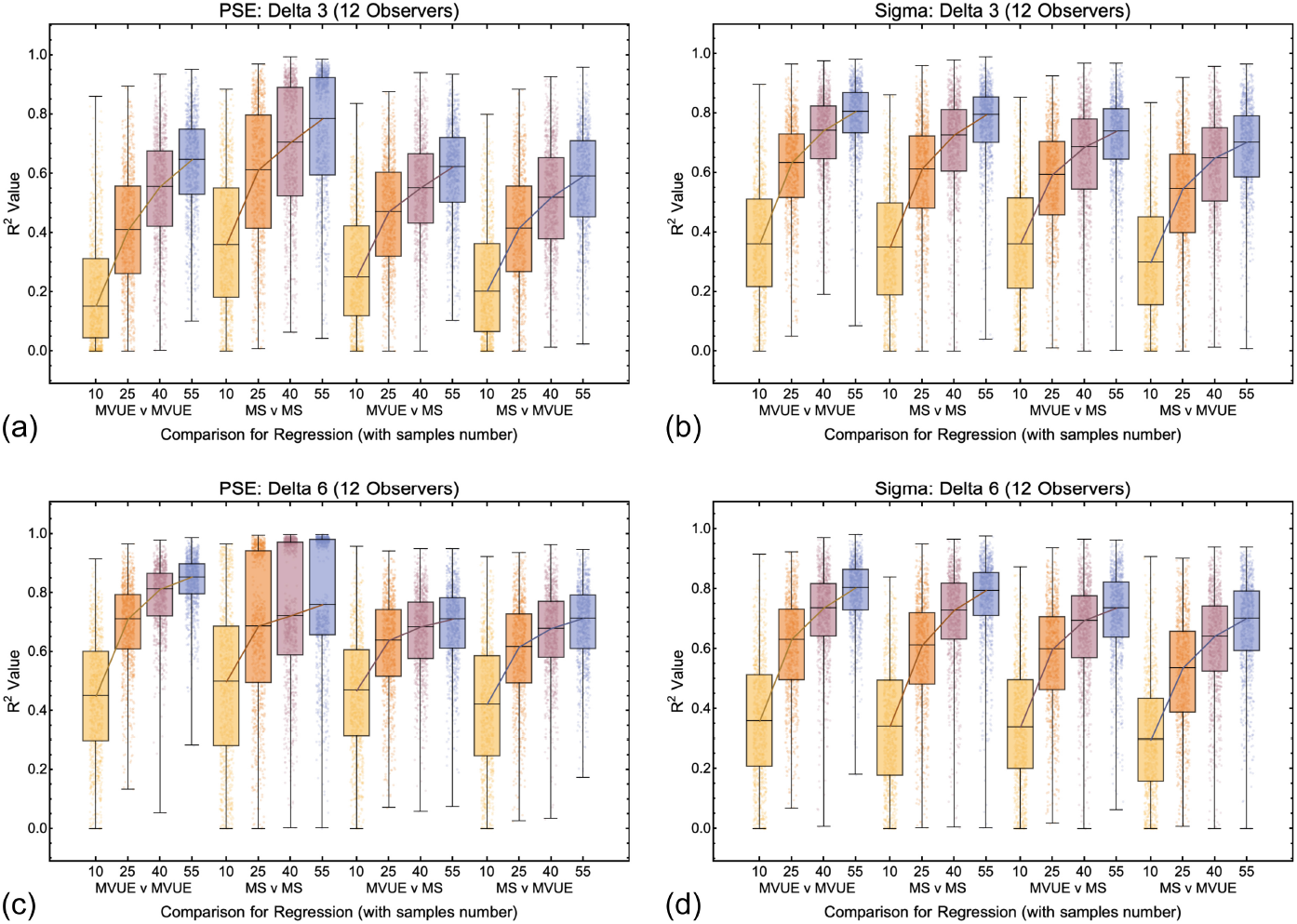
Box and whisker plots showing the distribution of R^2^ values for all conditions and comparisons in which there were 12 simulated observers per experiment for the 3mm (a and b) and 6mm (c and d) cue conflict (delta) conditions. The format is the same as Figure 9.

**Figure 11:**
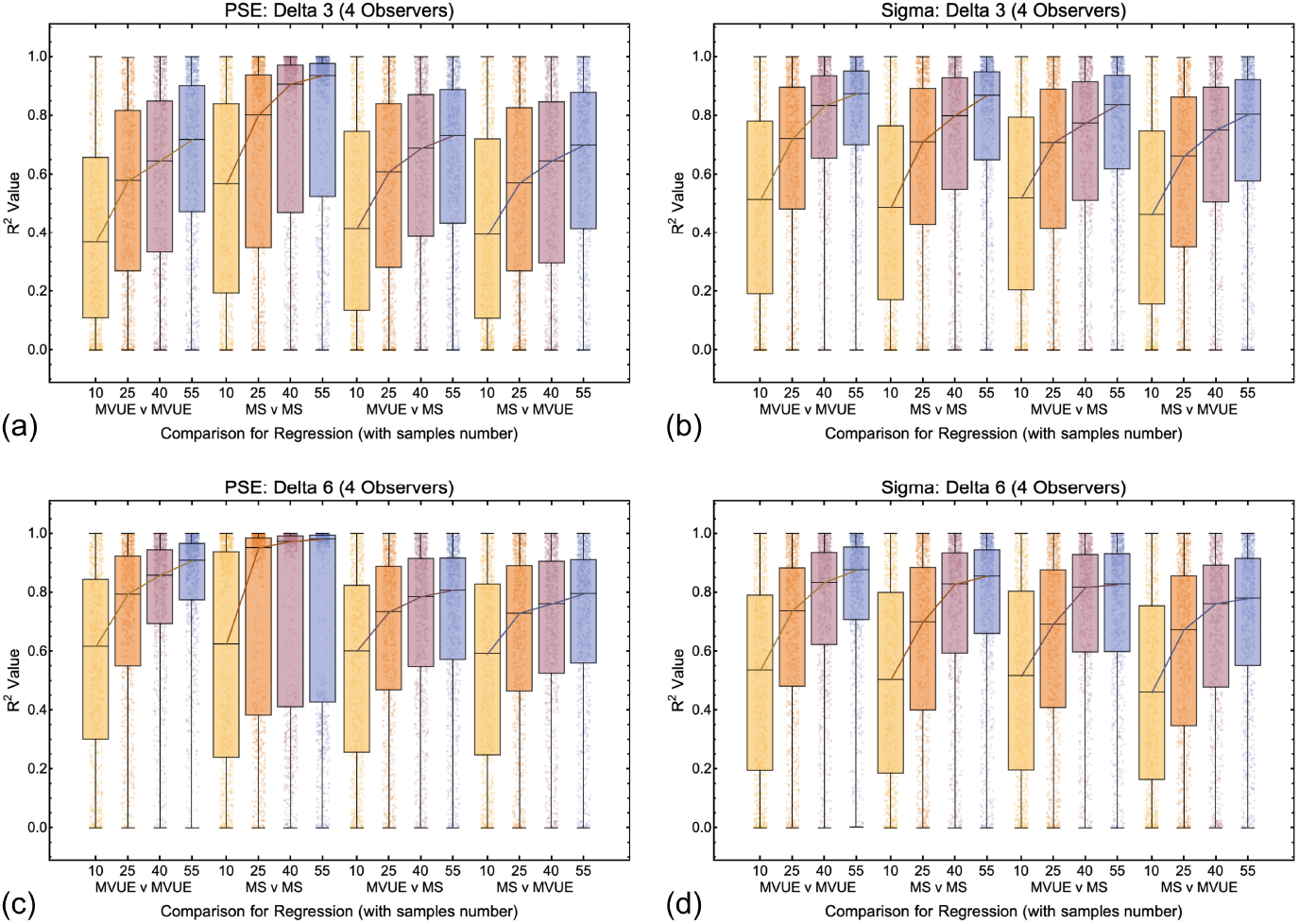
Box and whisker plots showing the distribution of R^2^ values for all conditions and comparisons in which there were 4 simulated observers per experiment for the 3mm (a and b) and 6mm (c and d) cue conflict (delta) conditions. The format is the same as Figures 10 and 11.

Figure 12 shows the overlap (Equation 10) for the distributions where a population of observers behaving in accordance with MVUE or MS were compared to the experimentally derived predictions of MVUE and MS. As expected, (1) the distribution overlap decreases with increasing amounts of data collected per function, (2) for the PSE distributions, the distribution overlap is less with a Δ of 6mm versus 3mm, and (3) the delta magnitude has no effect on the overlap of the sigma distributions. Problematically the distribution overlap is greater than 50% for virtually all conditions. This strongly questions one’s ability to use R^2^ to assess the extent to which a set of data are consistent with the predictions of MVUE. The precise amount of quantitative overlap acceptable for an experiment would be a judgement on the part of the experimenter.

**Figure 12:**
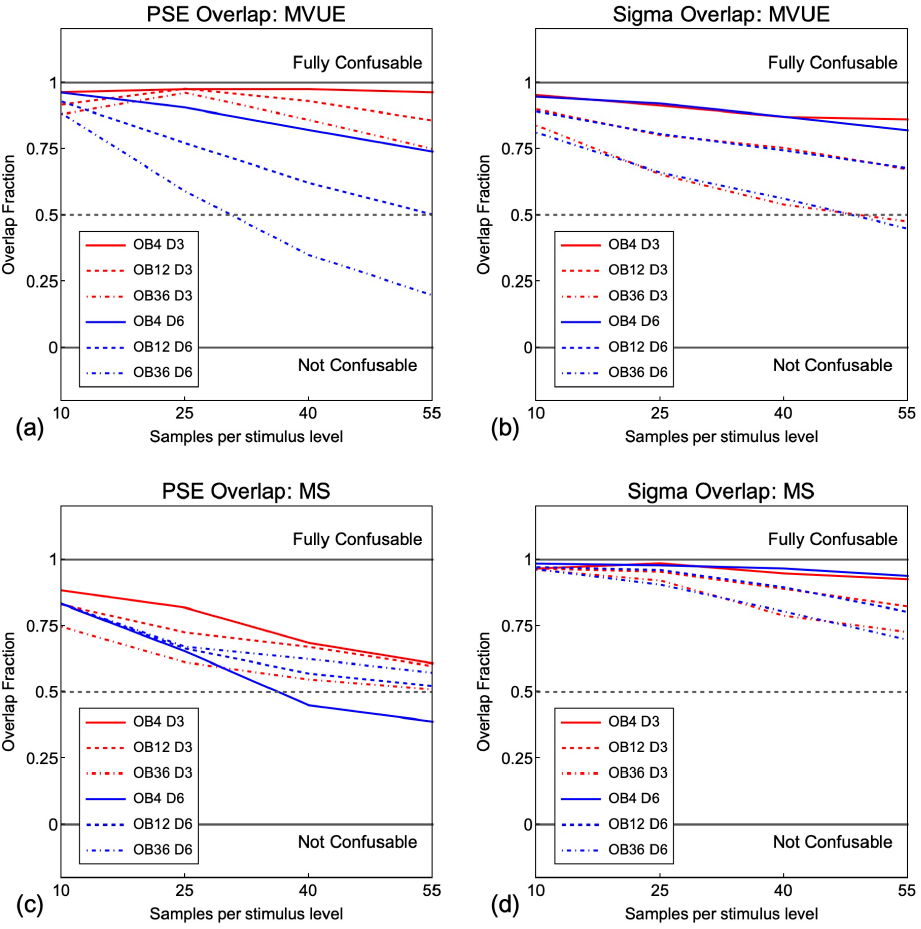
Overlap of the smooth kernel distributions of R^2^ value values produced from fitting a first order polynomial to observed data from observers behaving in accordance with MVUE against the experimentally derived predictions of MVUE and MS (PSE in (a) and Sigma in (b)), and a set of observers behaving in accordance with MS against the experimentally derived predictions of MVUE and MS (PSE in (c) and Sigma in (d)). Red lines are for the 3mm cu conflict and blue lines are for the 6mm cue conflict. “OB” is the number of observers in each simulated experiment and “D” the magnitude of the cue conflict in mm. An overlap value of 1 (upper solid grey line) means that the distributions (examples shown in Figure 8 e and f) completely overlap and are “fully confusable” and overlap of 0 means that the distributions do not overlap at all and are thus “not confusable”. The dashed grey line shows the case where the distributions overlap by 50%.

**Figure 13:**
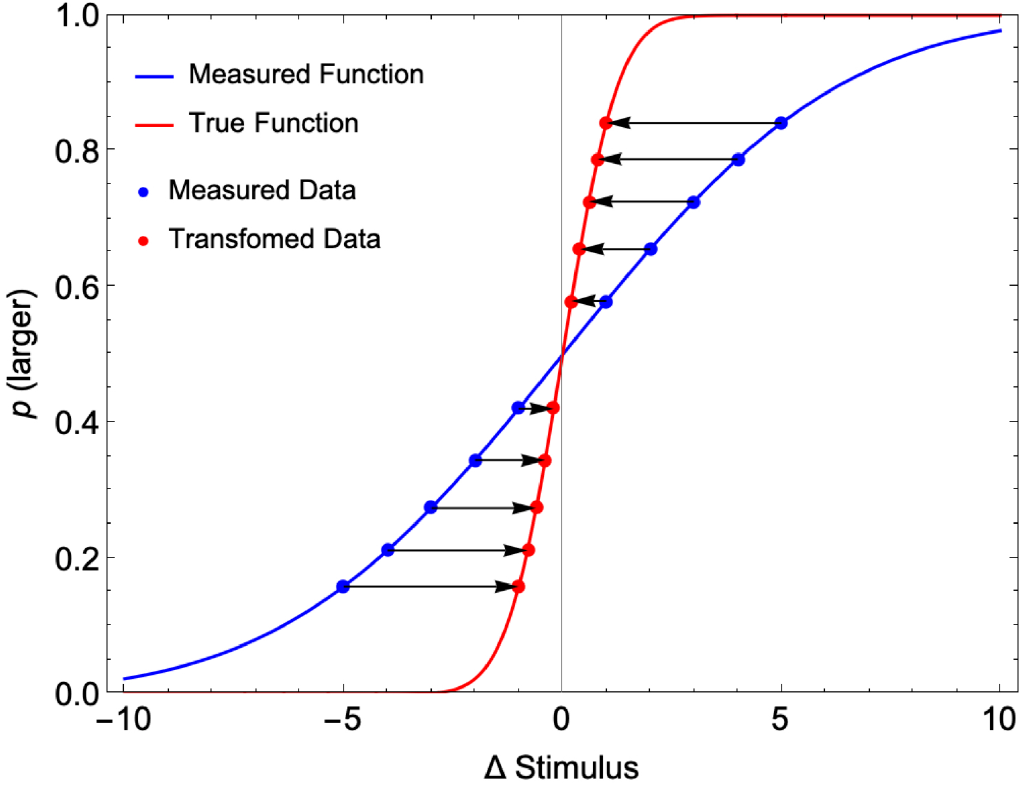
An experimenter presents a range of stimuli Δ S_A_(Δ Stimulus) and for each of these measures the probability of a “larger” response (“Measured Data”, shown as blue points). This is done in the presence of a conflicting cue, S_N_, which signals no change across intervals. For this visual example, σ_A_ = 1 and σ_N_ = 0.5, therefore w_A_ = 0.2 and w_N_ = 0.8 (Equations 3 and 4). The experimentally measured points are consistent with a measured psychometric function (blue Cumulative Gaussian function (given by Equation 31)). This function has a standard deviation 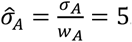. In reality, each stimulus ΔS_A_(i) is in fact cue conflict stimuli ΔS_C_(i) (given by Equation 25), thus the data should be shifted along the x-axis toward Δ Stimulus = 0 (black by arrows) to accurately plot the function. These shifted points (“Transformed Data”, shown as red points, Equation 26) are consistent with the true underlying psychometric function for the cue S_A_ (red Cumulative Gaussian function (given by Equation 30)). This function is steeper than the (measured) blue function because for a measured p(larger), the Δ Stimulus was in fact smaller than the experimenter had planned (due to the cue conflict).

An additional problem is that the *R*^2^ statistic that experimenters report *does not* measure the deviation of the data from the predictions of a cue integration model (even though it is often stated in this way), rather, the *R*^2^ statistic gives a measure of the fit *relative to the polynomial*. The predicted values of a cue integration model could be off by any arbitrary amount or have the *opposite* relationship between predictions and data, and experimenters could still obtain an *R*^2^ close to 1. Thus, a regression analysis negates one of the key benefits of MVUE (and other cue integration models), which is the ability to predict the absolute value of the integrated cues percept and its reliability and then compare this to that observed experimentally. Tests do exist to determine whether the intercept and slope differ from predicted model values, but these are rarely reported and are definitively not shown by the *R*^2^ statistic alone.

## Discussion

In any area of science, it is the job of a scientist to design experiments which can best distinguish between alternative models of the underlying phenomena. Unfortunately, in the area of cue integration this is rarely done. There are a wide range of competing models for how human observers might integrate information from sensory cues (Beierholm et al., 2009; Jones, 2016; Körding et al., 2007; Mamassian et al., 2002; Trommershauser et al., 2011), but in many instances the results of an experiment are simply visually inspected relative to the predictions of the experimenters preferred model (Negen et al., 2018; Rohde et al., 2016). This is problematic due to the small benefit accrued by models such as MVUE, the highly correlated predictions provided by alternative candidate models, and the fundamental misconceptions researchers have about how error bars relate to statistical significance (Belia et al., 2005; Cumming et al., 2007).

Whilst the numerous assumptions the MVUE model (and others) are known, these are rarely tested, instead experimenters typically assume that the assumptions are met and claim support for MVUE, often in the absence of statistical analysis and/or sufficient model comparison. The present paper aimed to draw attention to the assumptions of MVUE and to introduce a principled method by which to determine the probability with which a population of observers behaving in accordance with one model of cue integration can be experimentally distinguished from the predictions of alternative models. This showed that the experimental approach taken in many studies results in a poor ability to distinguish between alternative models (and thus the claim support for MVUE). At all decision points the simulations were designed to be (1) consistent with published guidelines stating how to test models of cue integration (Rohde et al., 2016), (2) consistent with the existing literature (Ernst & Banks, 2002), and (3) consistent with best practice as regards experimental methods (Fründ et al., 2011; Kingdom & Prins, 2016; Prins, 2012, 2013; Rohde et al., 2016; Wichmann & Hill, 2001a, 2001b).

Additionally, many of the nuisance parameters which would impede an experimenter’s ability to distinguish between models were not simulated. For example, for our simulated observers there was (1) statistical independence between trials, with no learning or boredom effects (Fründ et al., 2011), (2) a known generative function underlying behaviour (Kingdom & Prins, 2016; Murray & Morgenstern, 2010), (3) no perceptual bias (Scarfe & Hibbard, 2011), (4) stimulus values for the psychometric function were centred on the true mean of the psychometric function, (5) simulated observers exhibited no lapses (Prins, 2012; Wichmann & Hill, 2001a, 2001b), (6) the simulated data were not contaminated by the effect of decisional (or other sources of) noise (Hillis et al., 2004), (7) cues were statistically independent from one another (Oruc et al., 2003) and (8) there were no conflicting sources of sensory information (Watt et al., 2005). As a result, the simulations presented are highly likely to *overestimate* one’s ability to experimentally distinguish between models. These nuisance factors are known problems across all types of behavioural experiment, so are likely to be present to some extent in most studies. As a result, experimenters have designed experiments to eliminate them as best as possible, or where this is not possible, examined the effect they could have on the data (see Hillis et al., 2004; Oruc et al., 2003; Watt et al., 2005)

### Controlling for the effects of conflicting cues when measuring “single cue” sensitivities

A grounding assumption of the cue integration literature is that there exist separable sources of sensory information which provide independent perceptual estimates about properties of the world (Ernst & Bulthoff, 2004). In practice, it rapidly becomes apparent just how difficult it is to experimentally isolate sensory cues and to eliminate alternate cues which are not of interest (Watt et al., 2005; Zabulis & Backus, 2004). In many instances it remains possible that observers are utilising sensory cues that the experimenter was not intending to be available (Ho et al., 2006; Saunders & Chen, 2015; Todd, 2015; Todd et al., 2010). Even more problematically, some experiments measure “single cue” sensitivities in the presence of a known conflicting sensory cue held constant (Murphy et al., 2013; Svarverud et al., 2010). Here we examine the consequences of this.

Let’s assume that an experimenter is using a two-interval forced choice experiment to measure the sensitivity of a cue *S*_*A*_ for judgements of size. On each trial, in one interval the experiment presents a “standard” stimulus and in the other interval a “comparison” stimulus, the difference between these being Δ*S*_*A*_. The observer must signal in which interval the “larger” stimulus was presented. Next, let’s assume that this is done in the presence of a conflicting “nuisance” cue, Δ*S*_*N*_, which is constant and signals that the stimulus is unchanged across intervals. This means that the “single cue” stimulus is in fact an integrated cues stimulus and can be described as

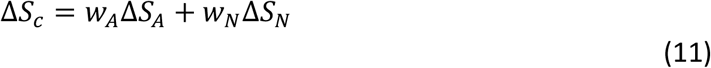

For each stimulus value Δ*S*_*c*_ (*i*), the experimenter measures *p*(“*larger”* | Δ*S*_*c*_(*i*)) and (with the assumption that the “standard” and “comparison” stimuli can be represented by Gaussian probability density functions) maps out a psychometric function by plotting *p*(*“larger”* | Δ*S*_*c*_ (*i*)) against Δ*S*_*A*_(*i*), then fits a Cumulative Gaussian to the data (blue data and function in Figure 11). Clearly, the experimenter will incorrectly estimate *σ*_*A*_ from this fitted function. More specifically, they will overestimate *σ*_*A*_ because each stimulus that they present is in fact an attenuated version of that which they intended (i.e., Δ*S*_*c*_ (*i*) < Δ*S*_*A*_(*i*)). The extent to which the experimenter misestimates *σ*_*A*_ will be a function of *w*_*N*_ (the weight given to the nuisance cue *S*_*N*_, which is signally no change across intervals). As *σ*_*N*_ → ∞, the weight given to the nuisance cue will approach zero (*w*_*N*_ → 0) and *σ*_*A*_ will be estimated accurately. However, for any non-infinite value of *σ*_*N*_, the experimenter will misestimate *σ*_*A*_.

In effect, what one needs to do is “warp” the x-axis of the measured psychometric function such that one is plotting *p*(*“larger”*) against Δ*S*_*c*_ (*i*) instead of Δ*S*_*A*_(*i*) (red data and function in Figure 11). To determine this “warping”, we can ask, what scale factor, *k*, would we need to apply to Δ*S*_*A*_ such that in all cases Δ*S*_*c*_ = Δ*S*_*A*_. Given *w*_*N*_ = 1 − *w*_*A*_, we can write this as

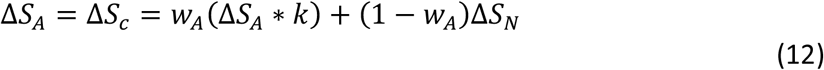

Recognising that Δ*S*_*N*_ = 0 and solving for *k*, we get

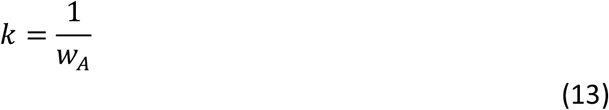

Intuitively we can see that this makes sense, as when *w*_*A*_ = 1, no scaling is required to combat the attenuation caused by Δ*S*_*N*_, because it receives zero weight, however, as soon as *w*_*A*_ < 1, scaling is needed (i.e., *k* > 1). Next, we can ask, given the true value of *σ*_*A*_, what would be our estimate, 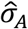, of this be in the presence of the conflicting nuisance cue. To do this we recognise that for a probability density function of a random variable *X* distributed according to *F*_*X*_(*x*), the probability density function of a variable *Y* = *g*(*X*) is also a random variable. If *g* is differentiable and *g*: ℝ → ℝ is a monotonic function, we can then use a *change of variables* to transform between probability density functions.

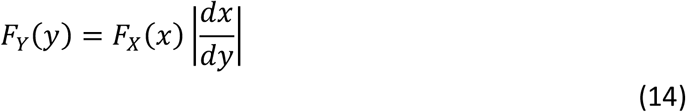

Here, (*x*) = *g*^−1^(*y*) and the support of *Y* is *g*(*x*) with the support of *X* being *x* (Blitzstein & Hwang, 2015). For our example, the Gaussian probability density function representing our cue *S*_*A*_ (red function in Figure 11) can be written as

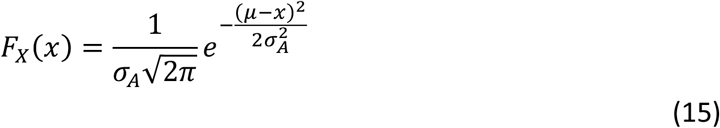

This function has a mean of *μ* and standard deviation of *σ*_*A*_. From Equation 13, using the transform *x* ∗ *k*, a change of variables gives

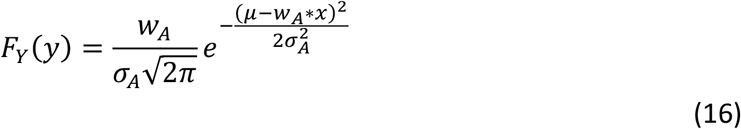

This represents our experimentally inferred probability density function for cue *S*_*A*_ (blue function in Figure 11). The standard deviation of *F*_*Y*_(*y*) is given by

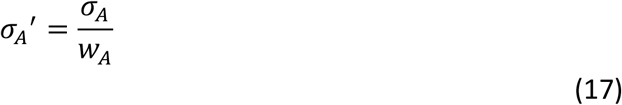

When weighting is given to the nuisance cue, *w*_*A*_ < 1, we overestimate the sigma of the underlying estimator, *σ*_*A*_*′*>*σ*_*A*_.

We can now determine the consequences this has for measuring the *relative reliability* of cues, which is the key variable needed for testing MVUE. Let’s say we have two cues *S*_*A*_ and *S*_*B*_ with standard deviations of *σ*_*A*_ and *σ*_*B*_ signalling a property of interest, *S*. We measure “single cue” sensitivity functions for each cue whilst holding the other cue constant. Because 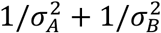 is a constant, *c*, the weights given to each cue are 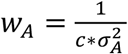 and 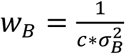, and given Equation 17, our experimental *estimates* of the true underlying standard deviations are given by 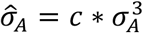 and 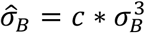. These are each larger than the true underlying values as they have been measured in the presence of a cue signally no change (Figure 11). The ratio of these estimates is given by

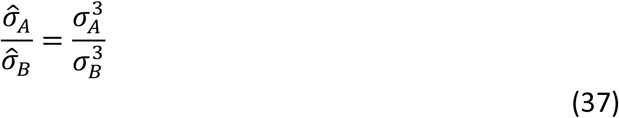

Thus, the ratio of the underlying sigma’s, which is the property we wish to estimate, is given by

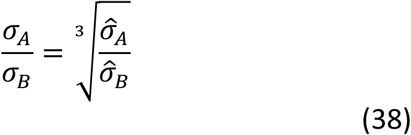

Therefore, if we infer from our experiment that *σ*_*A*_/*σ*_*B*_ = 1/27 the true sigma ratio is in fact 1/3 and we experimentally misestimate *σ*_*A*_/*σ*_*B*_ by a factor of ∼ 9. Studies which have measured the reliability of cues in the presence of a secondarily constant conflicting cue, (e.g. Murphy et al. (2013) and Svarverud et al. (2010)), will therefore have significantly overestimated the true cue relative reliabilities. As such, the data in these studies cannot be used to accurately test MVUE, without some form of correction. This analysis shows the critical importance of being able to isolate singles cues satisfactorily, or if one is not able to, correct for their influence when inferring relative cue reliabilities.

## Conclusion

The simplicity of the MVUE equations for cue integration is deceptive. A model’s simplicity is generally correlated with the number of assumptions it makes about the underlying phenomena. With more assumptions and a simpler model, there is a greater chance that the assumptions of the model will not be met. This will impact an experimenter’s ability to accurately test the predictions of the model. Even if one can be satisfied that the assumptions of MVUE hold in an experimental situation, MVUE provides correlated predictions with many other cue integration models (Arnold et al., 2019; Beierholm et al., 2009; Jones, 2016). Here we considered two such models, MS and PCS. It was shown that even when adopting published criteria describing how to best test the predictions of MVUE (Rohde et al., 2016), it could be very difficult to experimentally disambiguate between MVUE, MS and PCS. The analysis presented is only scratching the surface, as there are many ways in which sensory cues could be integrated (Jones, 2016), some of which may be even more difficult to disambiguate from MVUE.

Many studies claiming to support MVUE fail to consider alternative models satisfactorily, sample areas of the parameter space which poorly distinguish between competing models, and provide no statistical analysis related to the fit of MVUE to the data, or the relative fit of other alternative models. This questions the ability of these studies to conclude that sensory cues are integrated in accordance with MVUE. Whilst one could interpret the results presented here in a pessimistic fashion, the opposite is true. The results show clear, simple, and computationally attainable ways in which experimenters can correctly measure the variables needed to test models of cue integration and determine the probability with which a population of observers behaving in accordance with one model of sensory cue integration can be experimentally distinguished from the predictions of alternative models. Furthermore, it can be argued that the focus should not be upon attempting to prove that cues are integrated “optimally” based upon some criterion, but rather to simply focus on the factors that influence how cues are integrated (Rosas & Wichmann, 2011).

## Acknowledgements

The seeds of this project arose in the authors co-supervision, with Prof. Andrew Glennerster, of the PhD of Dr. Mark Adams. Preliminary results were presented at the Vision Sciences Society (VSS) 2016 meeting in St Pete’s beach, Florida (Scarfe and Glennerster (2016)). Feedback at VSS from Prof. Marc Ernst, Prof. Mike Landy, Prof. Jenny Read, Prof. Paul Hibbard, Prof. Loes van Dam, Prof. Andrew Glennerster, Dr. Katie Gray and other attendees helped convince the author that the project was of worthwhile interest to others. Prof. Andrew Glennerster and Dr. Katie Gray have provided advice and support through the project. I would also like to thank two anonymous reviewers whose comments greatly improved the manuscript.

## S1: Recording data from Ernst and Banks (2002)

Single cues sensitivities used for the simulations reported were estimated from Figure 3d of Ernst and Banks (2002). In order to gain estimates of the single cue sensitivities we viewed Figure 3d (as a pdf file) on a 4K computer monitor, so that the graph filled the majority of the screen. We then took the pixel coordinates of (1) the data points, (2) the minimum and maximum error bar position for each data-point and (3) the minimum and maximum values on the *x*- and *y*-axes. We were then able to compute the relative position of each data point (and error bar) in pixel coordinates on the *x*- and *y*-axis and covert these to the units shown in the graph by using the measured correspondence between pixel coordinates and axis units. Visual comparison of Figure 3 of the present paper and Figure 3d of Ernst and Banks shows that close correspondence achieved.

There was some inconsistently in Ernst and Banks (2002) as to how a “threshold” or “discrimination threshold” was defined. On page 430 the authors state, “The discrimination threshold is defined as the difference between the point of subjective equality (PSE) and the height of the comparison stimulus when it is judged taller than the standard stimulus 84% of the time”. However, on page 431 the authors state “… *T*_*H*_ and *T*_*V*_ are the haptic and visual thresholds (84% points in Fig. 3a)”. It is the first definition which is consistent with the mathematics i.e. the difference between the PSE and 84% point of the function being equal to the sigma of the fitted Cumulative Gaussian function.

Therefore, we cross checked our thresholds estimated from Figure 3d, with the thresholds calculated from the integrated cues functions in Figure 3b. Thresholds from Figure 3b were taken to be the difference between the point of subjective equality (PSE) and the 84% point on the function. When compared to the thresholds estimated from Figure 3d the difference in estimates was very small (average across data points of 0.23). We were therefore happy that definition of threshold was that of page 430 and that we had accurately estimated the thresholds and understood their relationship to the properties of the psychometric functions reported in the paper. Note: that for the purposes of the present paper all that was needed is an approximation of the exact values.

## S2: Example functions and goodness of fit

Figure S2a shows the true underlying functions for the minimum, maximum and base (middle) sigma values used in the current study as well as the stimulus levels at which the functions were sampled. As can be seen, consistent with Ernst and Banks (2002) Figure 3a, all functions straddle high and low performance levels needed for well fit functions (Wichmann & Hill, 2001a, 2001b). Figures S2b-e show examples of how these functions were sampled with our four sampling regimes (10, 25, 40 and 55 trials per stimulus level), with the maximum likelihood best fit functions and goodness of fit (see below) values shown in the legend. We have only shown these for just the Δ = 0 case, as for all delta values used the sampling range was shifted so as to be centred on the true mean of the underlying function. As is clear, for all sampling regimes the data are well fit by the Cumulative Gaussian functions.

**Figure S2:**
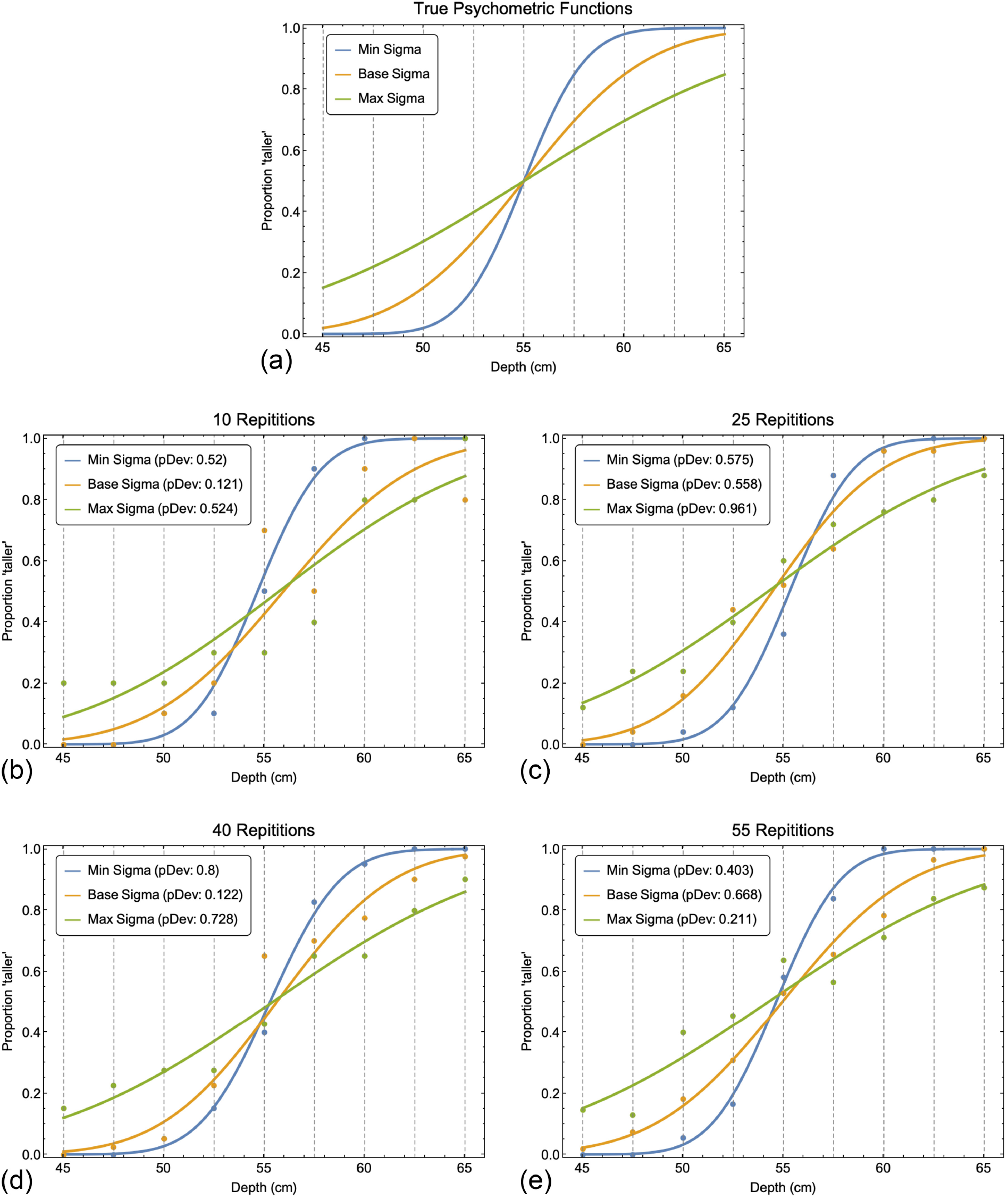
(a) shows the underlying “true” psychometric functions for the minimum, maximum and base (middle) sigma values used in the paper. The dashed vertical grey lines show the nine stimulus values at which these functions were sampled. (b) through (e) show examples of how the functions were sampled through simulation and fit with psychometric functions for the four data collection regimes used throughout the paper (10, 25, 40, and 55 repetitions per stimulus level). Inset in each graph is the goodness of fit value, pDev. This represents the probability with which the experimental data produced a higher likelihood ratio than that of the stimulated experiments based upon the target model. If this is greater than 0.05, the function is considered to fit the data well. See accompanying text for details.

Within the cue integration literature, the goodness of fit of a function and the criteria upon which a fit is considered unacceptable is rarely if ever stated (for example Ernst & Banks, 2002; Helbig & Ernst, 2007; Hillis et al., 2002). Thus, it is impossible to tell if a goodness of fit test was performed, and if one was, which test which was used, and the criteria adopted for rejecting a fitted function. Given that the fit of data to the MVUE model is normally assessed by eye, it is likely that this is also the case for the fit of individual psychometric functions (Kingdom & Prins, 2016). The Palamedes toolbox (Prins & Kingdom, 2009) used in the present study implements a bootstrapped likelihood ratio test to assess the goodness of fit of a psychometric function. The logic of the test is as follows (Kingdom & Prins, 2016).

As detailed in the main text, when fitting a psychometric function to some data the experimenter assumes: (1) the observer does not improve or degrade at the task they are performing over time, (2) each perceptual judgement an observer makes is statistically independent of all others, and (3) performance of the observer can be well characterised by the psychometric function that the experimenter is choosing to fit to the data. These assumptions combined can be referred to as the “target model”. The validity of the target model can be assessed by comparing it to a “saturated model” which only assumes (1) and (2). Thus, in the saturated model, the probability of response for one stimulus level is completely independent on the probability of response for any other stimulus level i.e. no psychometric function is assumed.

The target model is “nested” under the saturated model, as it is a single specific case of the saturated model. Thus, the likelihood associated with the fit of the target model can never produce a better fit than that of the less restrictive saturated model. For a given set of data one can calculate the likelihood ratio (likelihood of the target model / likelihood of the saturated model) which will, by definition, be less than or equal to 1. It will only be equal to one if the target and saturated models provide as good a fit as one another. The likelihood ratio test simulates a set of experiments through a bootstrap procedure and for each calculates the likelihood ratio. The probability with which the experimental data produces a higher likelihood ratio than that of the stimulated experiments is calculated (*pDev* in Figure S2). If this probability is less than 0.05% the goodness of fit is deemed poor. As with any *p*-value, the 0.05% cut-off is a completely arbitrary convention (Kingdom & Prins, 2016). Thus, some experimenters may adopt this and others not. This mirrors the open discussion about the use of *p*-values for general statistical analysis.

For the present study, it was computationally unfeasible to run a bootstrapped likelihood ratio test for each of the ∼15.3 million simulated functions (even when using MATLAB’s Parallel processing toolbox to spread the computational load over the 8-Core Intel Core i9 available to the author this would have taken ∼1-2 months of constant processing). Nevertheless, we wanted to assess the extent to which the maximum likelihood fit functions would in general be considered well fit. Therefore, for the maximum and minimum cue sigma value used in the paper (i.e. shallowest and steepest psychometric functions), we simulated data for 1000 observers, fit Cumulative Gaussian psychometric functions to the data (as described in the main text) and assessed the goodness of fit using the bootstrapped likelihood ratio test (1000 bootstraps). We did this for our four sampling regimes: 10, 25, 40 and 55 trials per stimulus level.

Based upon the 0.05% criteria for a cut-off between well and poorly fit function (*pDev* in Figure S2), virtually all functions would have been classed as well fit, regardless of data collection regime of the slope of the underlying function (Table T1; overall average 94.95%). As would be expected, this was true for all Delta levels. This is because the sampling range was always centred on the true mean of the function, so the values for Delta 0, 3 and 6 in Table T1 are effectively replications of one another. This confirms across 24000 fitted functions what can be seen in the example functions of Figure S2 i.e. that the data are well fit by the psychometric functions. We can therefore be satisfied that the around 94.95% of all functions reported in the paper would have been classed as well fit based on this criteria. See also the criteria adopted for rejecting psychometric functions discussed in the main body of the text.

**Table T2:** Shows the percentage of psychometric functions which would be classified as well fit based upon the bootstrapped likelihood ratio test described in the main text. The percentage of well fit functions is shown for the minimum and maximum sigma used in the simulations of the paper, and for each combination of trials per stimulus value on the psychometric function and cue conflict level (cue delta in mm).

| Sigma / Trials | Delta 0 | Delta 3 | Delta 6 |
| --- | --- | --- | --- |
| Min 10 | 93.8% | 95.5% | 95.4% |
| Max 10 | 95.5% | 95.2% | 95.5% |
| Min 25 | 96.2% | 94.5% | 94.6% |
| Max 25 | 95.6% | 96.7% | 95.4% |
| Min 40 | 95.7% | 95.5% | 94.1% |
| Max 40 | 94.7% | 95% | 95.2% |
| Min 55 | 95.5% | 94.1% | 94.4% |
| Max 55 | 94.7% | 93.7% | 94.7% |
| Mean Value | **94.91%** | **95.03%** | **94.91%** |

**Figure S3:**
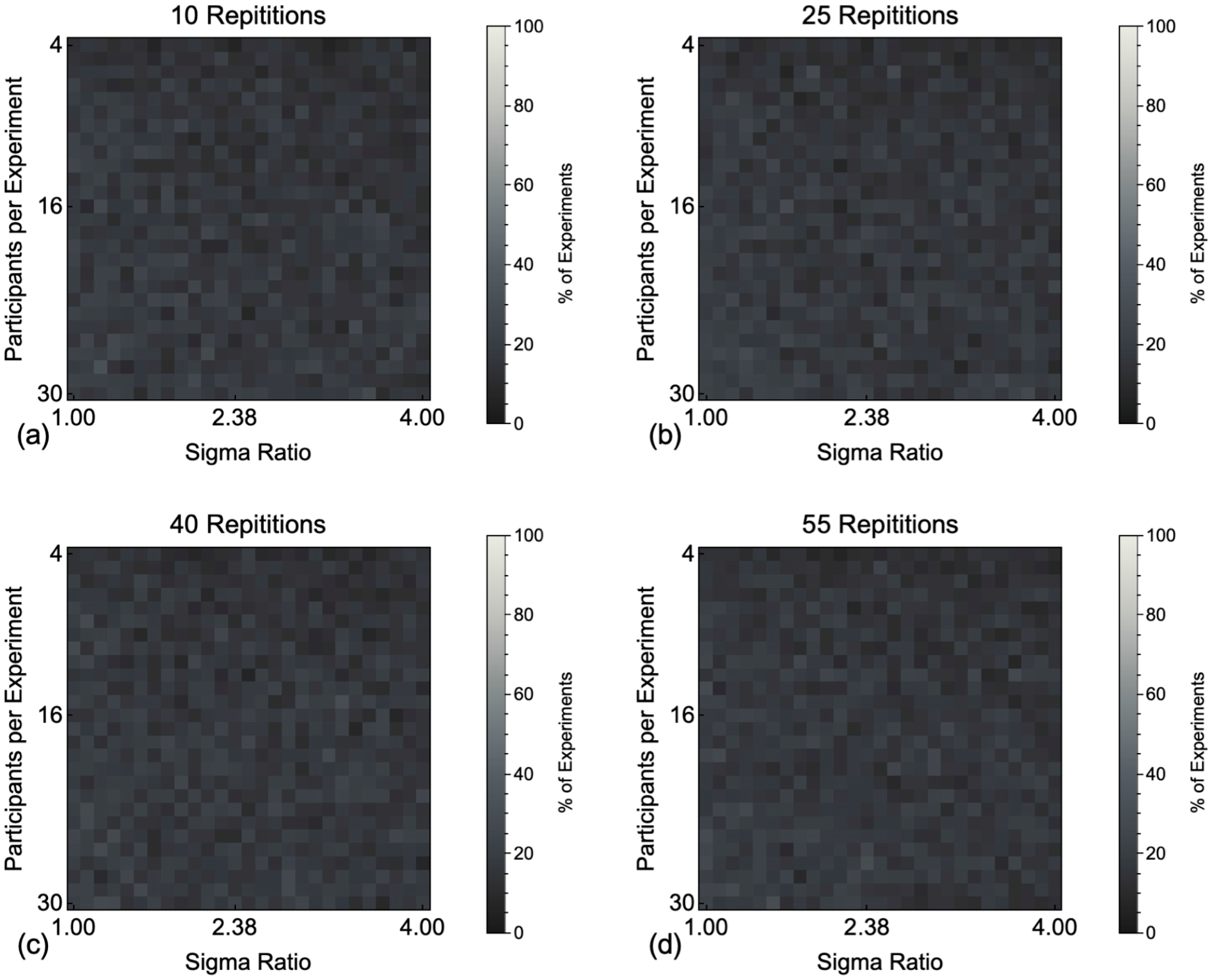
Shows the percentage of experiments in which the mean of the Cumulative Gaussian functions fit to our simulated population of MVUE observers could be statistically distinguished from the experimentally derived prediction of MS, when there is zero cue conflict. Each pixel in the image shows this percentage as calculated across 100 simulated experiments, of a given sigma ratio and number of participants. The four panes show this for (a) 10, (b) 25, (c) 40 and (d) 55, simulated trials per stimulus level on the psychometric function.

**Figure S4:**
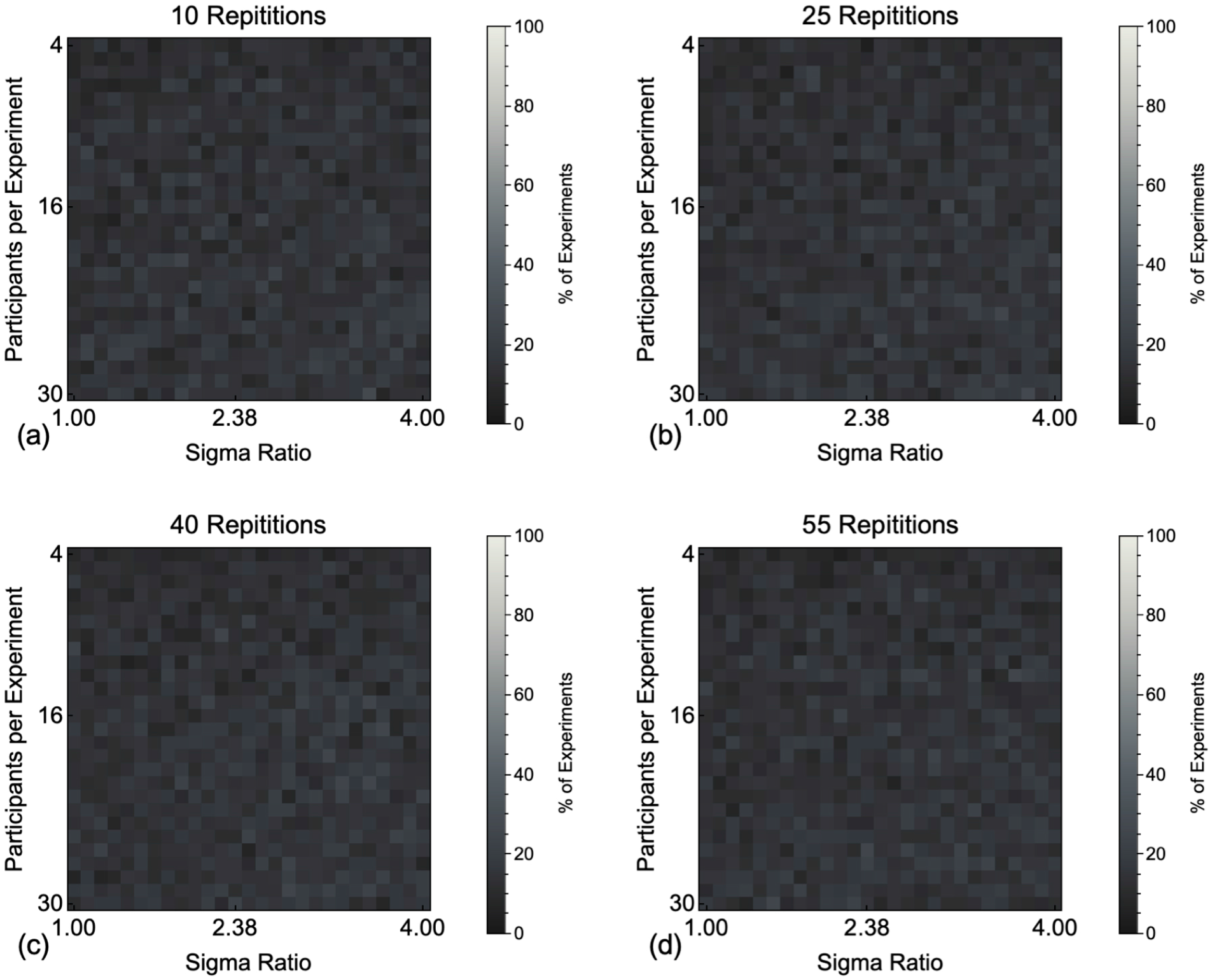
Shows the percentage of experiments in which the mean of the Cumulative Gaussian functions fit to our simulated population of MVUE observers could be statistically distinguished from the experimentally derived prediction of PCS, when there is zero cue conflict. Each pixel in the image shows this percentage as calculated across 100 simulated experiments, of a given sigma ratio and number of participants. The four panes show this for (a) 10, (b) 25, (c) 40 and (d) 55, simulated trials per stimulus level on the psychometric function.

**Figure S5:**
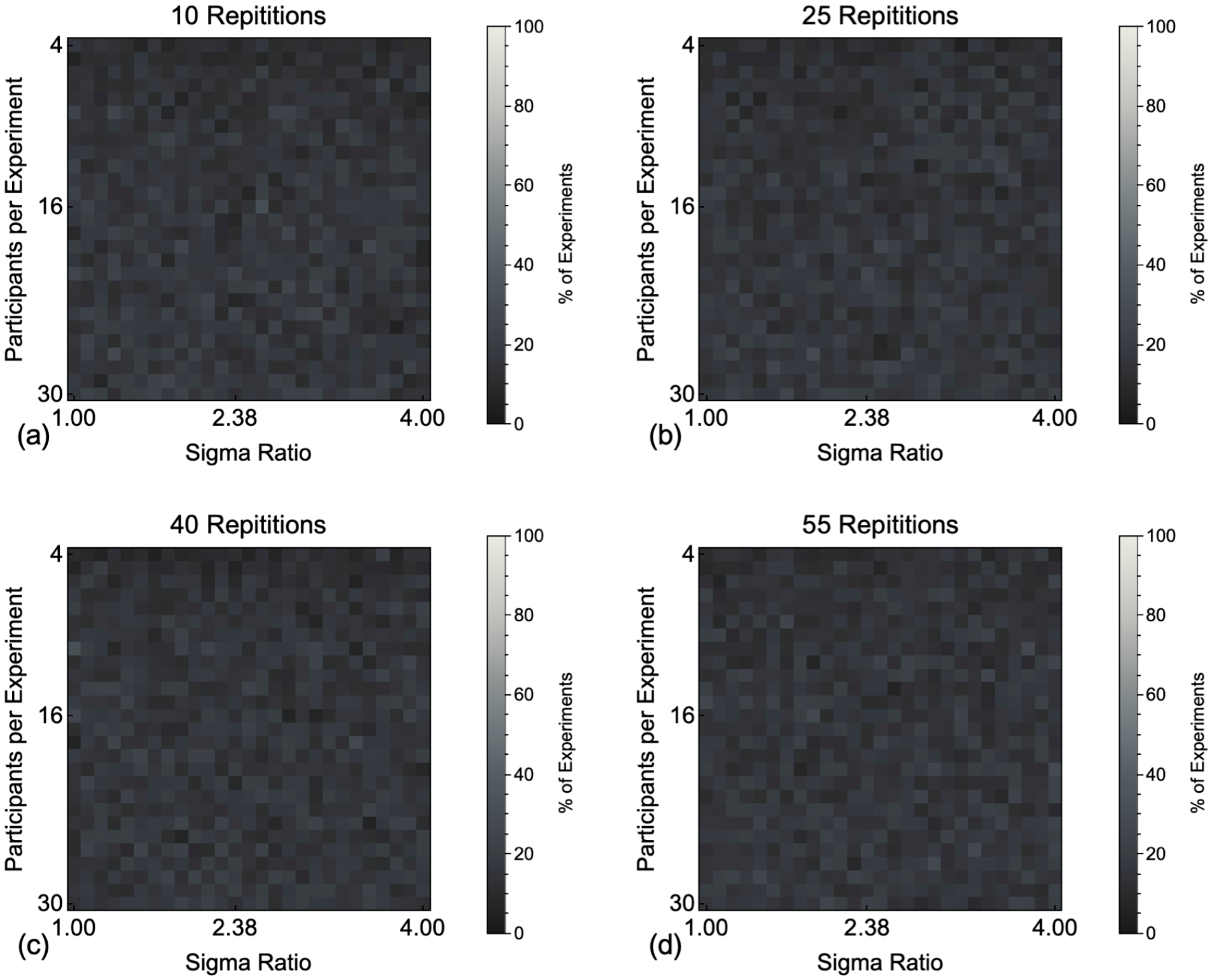
Shows the percentage of experiments in which the mean of the Cumulative Gaussian functions fit to our simulated population of MVUE observers could be statistically distinguished from the experimentally derived prediction of PCS with an experimental cue conflict of 3mm. Each pixel in the image shows this percentage as calculated across 100 simulated experiments, of a given sigma ratio and number of participants. The four panes show this for (a) 10, (b) 25, (c) 40 and (d) 55, simulated trials per stimulus level on the psychometric function.

**Figure S6:**
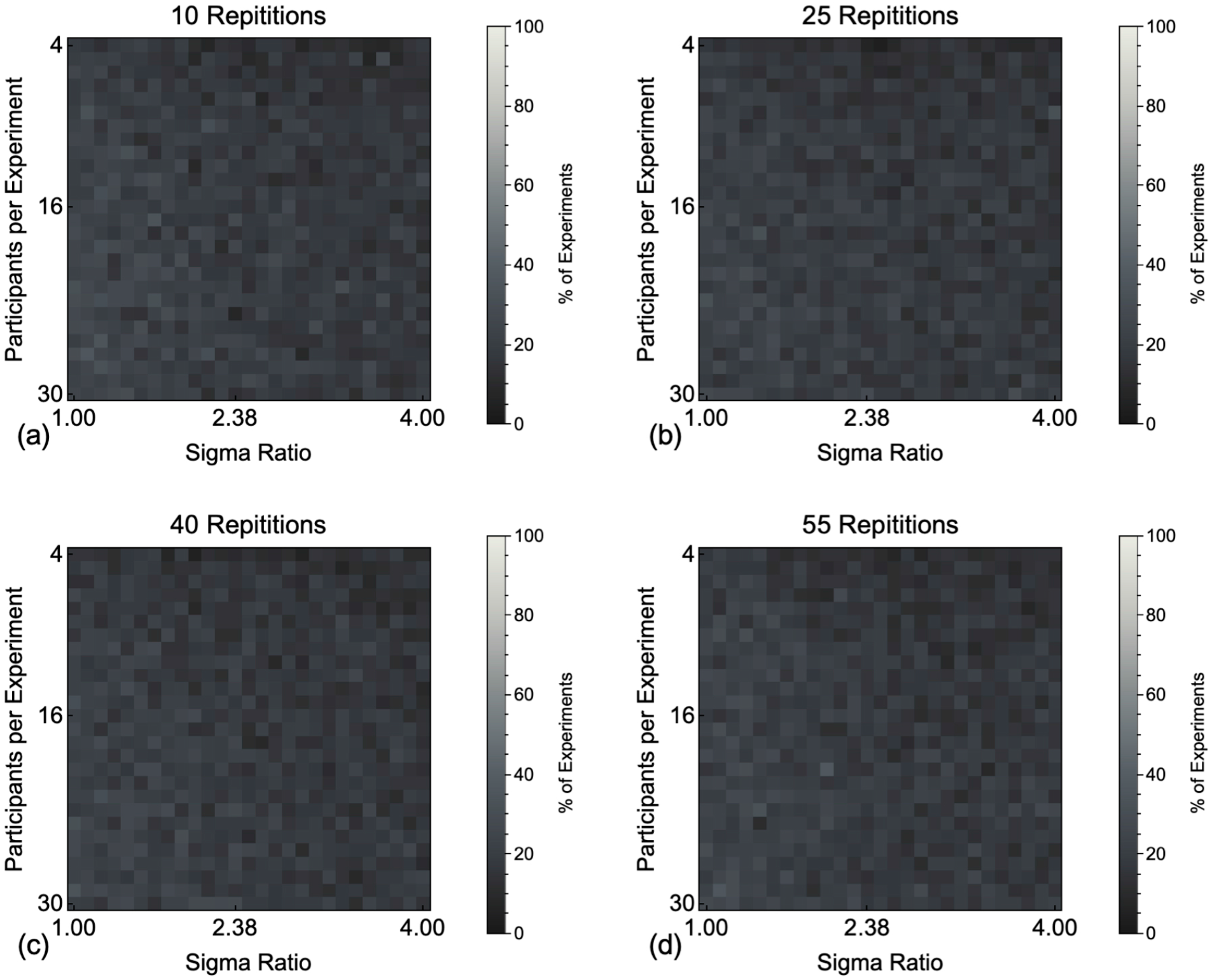
Shows the percentage of experiments in which the mean of the Cumulative Gaussian functions fit to our simulated population of MVUE observers could be statistically distinguished from the experimentally derived prediction of PCS with an experimental cue conflict of 6mm. Each pixel in the image shows this percentage as calculated across 100 simulated experiments, of a given sigma ratio and number of participants. The four panes show this for (a) 10, (b) 25, (c) 40 and (d) 55, simulated trials per stimulus level on the psychometric function.

## Notes

### Competing Interest Statement

The authors have declared no competing interest.

### Summary of Updates

Based upon the current review process the whole paper has been updated.

## References

Acerbi, L., Dokka, K., Angelaki, D. E., & Ma, W. J. (2018). Bayesian comparison of explicit and implicit causal inference strategies in multisensory heading perception. Plos Computational Biology, 14(7), e1006110. https://doi.org/10.1371/journal.pcbi.1006110

Adams, W. J., Banks, M. S., & van Ee, R. (2001). Adaptation to three-dimensional distortions in human vision. Nat Neurosci, 4(11), 1063–1064. https://doi.org/10.1038/nn729

Arnold, D. H., Petrie, K., Murray, C., & Johnston, A. (2019). Suboptimal human multisensory cue combination. Sci Rep, 9(1), 5155. https://doi.org/10.1038/s41598-018-37888-7

Beierholm, U., Shams, L., Körding, K., & Ma, W. J. (2009). Comparing Bayesian models for multisensory cue combination without mandatory fusion Advances in Neural Information Processing Systems 20. Advances in Neural Information Processing Systems,

Belia, S., Fidler, F., Williams, J., & Cumming, G. (2005). Researchers misunderstand confidence intervals and standard error bars. Psychol Methods, 10(4), 389–396. https://doi.org/10.1037/1082-989X.10.4.389

Blitzstein, J. K., & Hwang, J. (2015). Introduction to Probability. CRC Press.

Bradshaw, M. F., Parton, A. D., & Glennerster, A. (2000). The task-dependent use of binocular disparity and motion parallax information. Vision Research, 40(27), 3725–3734.

Burge, J., Fowlkes, C. C., & Banks, M. S. (2010). Natural-scene statistics predict how the figure-ground cue of convexity affects human depth perception. Journal of Neuroscience, 30(21), 7269–7280. https://doi.org/10.1523/JNEUROSCI.5551-09.2010

Burge, J., Girshick, A. R., & Banks, M. S. (2010). Visual-haptic adaptation is determined by relative reliability. Journal of Neuroscience, 30(22), 7714–7721. https://doi.org/10.1523/JNEUROSCI.6427-09.2010

Burge, J., Peterson, M. A., & Palmer, S. E. (2005). Ordinal configural cues combine with metric disparity in depth perception. J Vis, 5(6), 534–542. https://doi.org/10.1167/5.6.5

Byrne, P. A., & Henriques, D. Y. (2013). When more is less: increasing allocentric visual information can switch visual-proprioceptive combination from an optimal to sub-optimal process. Neuropsychologia, 51(1), 26–37. https://doi.org/10.1016/j.neuropsychologia.2012.10.008

Cochran, W. G. (1937). Problems arising in the analysis of a series of similar experiments. Journal of the Royal Statistical Society, 4, 102–118.

Cumming, G., Fidler, F., & Vaux, D. L. (2007). Error bars in experimental biology. J Cell Biol, 177(1), 7–11. https://doi.org/10.1083/jcb.200611141

de Winkel, K. N., Katliar, M., Diers, D., & Bulthoff, H. H. (2018). Causal Inference in the Perception of Verticality. Sci Rep, 8(1), 5483. https://doi.org/10.1038/s41598-018-23838-w

Domini, F., & Caudek, C. (2009). The intrinsic constraint model and Fechnerian sensory scaling [Comparative Study Research Support, U.S. Gov’t, Non-P.H.S.]. J Vis, 9(2), 25 21–15. https://doi.org/10.1167/9.2.25

Ernst, M. O. (2006). A Bayesian view on multimodal cue integration. In G. Knoblich, I. M. Thornton, M. Grosjean, & M. Shiffrar (Eds.), Human body perception from the inside out (pp. 105–131). Oxford University Press.

Ernst, M. O., & Banks, M. S. (2002). Humans integrate visual and haptic information in a statistically optimal fashion. Nature, 415(6870), 429–433. https://doi.org/10.1038/415429a

Ernst, M. O., & Bulthoff, H. H. (2004). Merging the senses into a robust percept. Trends Cogn Sci, 8(4), 162–169. https://doi.org/10.1016/j.tics.2004.02.002

Ernst, M. O., & Di Luca, M. (2011). Multisensory perception: from integration to remapping. In J. Trommershauser, K. P. Körding, & M. S. Landy (Eds.), Sensory Cue Integration (pp. 224–250). Oxford University Press.

Fischer, J., & Whitney, D. (2014). Serial dependence in visual perception. Nat Neurosci, 17(5), 738–743. https://doi.org/10.1038/nn.3689

Fründ, I., Haenel, N. V., & Wichmann, F. A. (2011). Inference for psychometric functions in the presence of nonstationary behavior. J Vis, 11(6). https://doi.org/10.1167/11.6.16

Gepshtein, S., Burge, J., Ernst, M. O., & Banks, M. S. (2005). The combination of vision and touch depends on spatial proximity. J Vis, 5(11), 1013–1023. https://doi.org/10.1167/5.11.7

Girshick, A. R., & Banks, M. S. (2009). Probabilistic combination of slant information: weighted averaging and robustness as optimal percepts. J Vis, 9(9), 8 1–20. https://doi.org/10.1167/9.9.8

Glennerster, A., Tcheang, L., Gilson, S. J., Fitzgibbon, A. W., & Parker, A. J. (2006). Humans ignore motion and stereo cues in favor of a fictional stable world. Curr Biol, 16(4), 428–432. https://doi.org/10.1016/j.cub.2006.01.019

Green, D. M., & Swets, J. A. (1974). Signal Detection Theory and Psychophysics. Cambridge University Press.

Helbig, H. B., & Ernst, M. O. (2007). Optimal integration of shape information from vision and touch. Exp Brain Res, 179(4), 595–606. https://doi.org/10.1007/s00221-006-0814-y

Henriques, D. Y., & Cressman, E. K. (2012). Visuomotor adaptation and proprioceptive recalibration. J Mot Behav, 44(6), 435–444. https://doi.org/10.1080/00222895.2012.659232

Hillis, J. M., Ernst, M. O., Banks, M. S., & Landy, M. S. (2002). Combining sensory information: Mandatory fusion within, but not between, senses. Science, 298(5598), 1627-1630. <Go to ISI>://000179361600051

Hillis, J. M., Watt, S. J., Landy, M. S., & Banks, M. S. (2004). Slant from texture and disparity cues: optimal cue combination. J Vis, 4(12), 967–992. https://doi.org/10.1167/4.12.1

Ho, Y. X., Landy, M. S., & Maloney, L. T. (2006). How direction of illumination affects visually perceived surface roughness. J Vis, 6(5), 634–648. https://doi.org/10.1167/6.5.8

Jacobs, R. A. (2002). What determines visual cue reliability? Trends in Cognitive Sciences, 6(8), 345-350. <Go to ISI>://000177263200009

Johnston, E. B., Cumming, B. G., & Landy, M. S. (1994). Integration of stereopsis and motion shape cues. Vision Res, 34(17), 2259–2275. https://www.ncbi.nlm.nih.gov/pubmed/7941420

Johnston, E. B., Cumming, B. G., & Parker, A. J. (1993). Integration of depth modules: stereopsis and texture. Vision Res, 33(5-6), 813-826. https://doi.org/Doi10.1016/0042-6989(93)90200-G

Jones, P. R. (2016). A tutorial on cue combination and Signal Detection Theory: Using changes in sensitivity to evaluate how observers integrate sensory information. Journal of Mathematical Psychology, 73, 117–139.

Kingdom, F. A. A., & Prins, N. (2010). Psychophysics: A Practical Introduction. (1st ed.). Academic Press.

Kingdom, F. A. A., & Prins, N. (2016). Psychophysics: A Practical Introduction. (2nd ed.). Academic Press.

Kiyonaga, A., Scimeca, J. M., Bliss, D. P., & Whitney, D. (2017). Serial Dependence across Perception, Attention, and Memory. Trends Cogn Sci, 21(7), 493–497. https://doi.org/10.1016/j.tics.2017.04.011

Knill, D. C., & Richards, W. (1996). Perception as Bayesian Inference. Cambridge University Press.

Knill, D. C., & Saunders, J. A. (2003). Do humans optimally integrate stereo and texture information for judgments of surface slant? Vision Research, 43(24), 2539–2558. <Go to ISI>://000185577700006

Koenderink, J. J., van Doorn, A. J., Kappers, A. M., & Lappin, J. S. (2002). Large-scale visual frontoparallels under full-cue conditions. Perception, 31(12), 1467–1475. https://doi.org/10.1068/p3295

Koenderink, J. J., van Doorn, A. J., Kappers, A. M., & Todd, J. T. (2002). Pappus in optical space. Percept Psychophys, 64(3), 380–391. https://www.ncbi.nlm.nih.gov/pubmed/12049279

Koenderink, J. J., van Doorn, A. J., & Lappin, J. S. (2000). Direct measurement of the curvature of visual space. Perception, 29(1), 69–79. https://doi.org/10.1068/p2921

Kontsevich, L. L., & Tyler, C. W. (1999). Bayesian adaptive estimation of psychometric slope and threshold. Vision Res, 39(16), 2729–2737. https://doi.org/Doi10.1016/S0042-6989(98)00285-5

Körding, K. P., Beierholm, U., Ma, W. J., Quartz, S., Tenenbaum, J. B., & Shams, L. (2007). Causal inference in multisensory perception [Clinical Trial Research Support, Non-U.S. Gov’t]. PloS one, 2(9), e943. https://doi.org/10.1371/journal.pone.0000943

Kruschke, J. K. (2010). What to believe: Bayesian methods for data analysis. Trends Cogn Sci, 14(7), 293–300. https://doi.org/10.1016/j.tics.2010.05.001

Kruschke, J. K. (2011). Doing Bayesian Data Analysis. Elsevier.

Kuss, M., Jakel, F., & Wichmann, F. A. (2005). Bayesian inference for psychometric functions. J Vis, 5(5), 478–492. https://doi.org/10.1167/5.5.8

Lages, M., & Jaworska, K. (2012). How Predictable are “Spontaneous Decisions” and “Hidden Intentions”? Comparing Classification Results Based on Previous Responses with Multivariate Pattern Analysis of fMRI BOLD Signals. Front Psychol, 3, 56. https://doi.org/10.3389/fpsyg.2012.00056

Landy, M. S., Maloney, L. T., Johnston, E. B., & Young, M. (1995). Measurement and modeling of depth cue combination: in defense of weak fusion. Vision Res, 35(3), 389–412. https://www.ncbi.nlm.nih.gov/pubmed/7892735

Leek, M. R. (2001). Adaptive procedures in psychophysical research. Percept Psychophys, 63(8), 1279–1292. https://doi.org/10.3758/bf03194543

Liberman, A., Fischer, J., & Whitney, D. (2014). Serial dependence in the perception of faces. Curr Biol, 24(21), 2569–2574. https://doi.org/10.1016/j.cub.2014.09.025

Liberman, A., Manassi, M., & Whitney, D. (2018). Serial dependence promotes the stability of perceived emotional expression depending on face similarity. Atten Percept Psychophys, 80(6), 1461–1473. https://doi.org/10.3758/s13414-018-1533-8

Liberman, A., Zhang, K., & Whitney, D. (2016). Serial dependence promotes object stability during occlusion. J Vis, 16(15), 16. https://doi.org/10.1167/16.15.16

Lovell, P. G., Bloj, M., & Harris, J. M. (2012). Optimal integration of shading and binocular disparity for depth perception. J Vis, 12(1). https://doi.org/10.1167/12.1.1

Mamassian, P., Landy, M. S., & Maloney, L. T. (2002). Bayesian Modelling of Visual Perception. In R. P. N. Rao, B. A. Olshausen, & M. S. Lewicki (Eds.), Probabilistic Models of the Brain: Perception and Neural Function (pp. 13–36).

McLaughlin, S. C., & Webster, R. G. (1967). Changes in the straight-ahead eye position during adaptation to wedge prisms. Percept Psychophysics, 2(1), 37–44.

Murphy, A. P., Ban, H., & Welchman, A. E. (2013). Integration of texture and disparity cues to surface slant in dorsal visual cortex. J Neurophysiol, 110(1), 190–203. https://doi.org/10.1152/jn.01055.2012

Murray, R. F., & Morgenstern, Y. (2010). Cue combination on the circle and the sphere. J Vis, 10(11), 15. https://doi.org/10.1167/10.11.15

Nardini, M., Jones, P., Bedford, R., & Braddick, O. (2008). Development of cue integration in human navigation. Curr Biol, 18(9), 689–693. https://doi.org/10.1016/j.cub.2008.04.021

Negen, J., Wen, L., Thaler, L., & Nardini, M. (2018). Bayes-Like Integration of a New Sensory Skill with Vision. Sci Rep, 8(1), 16880. https://doi.org/10.1038/s41598-018-35046-7

Oruc, I., Maloney, L. T., & Landy, M. S. (2003). Weighted linear cue combination with possibly correlated error. Vision Res, 43(23), 2451–2468. https://www.ncbi.nlm.nih.gov/pubmed/12972395

Pastore, M., & Calcagni, A. (2019). Measuring Distribution Similarities Between Samples: A Distribution-Free Overlapping Index. Front Psychol, 10, 1089. https://doi.org/10.3389/fpsyg.2019.01089

Pentland, A. (1980). Maximum likelihood estimation: the best PEST. Percept Psychophys, 28(4), 377–379. https://doi.org/10.3758/bf03204398

Prins, N. (2012). The psychometric function: the lapse rate revisited. J Vis, 12(6). https://doi.org/10.1167/12.6.25

Prins, N. (2013). The psi-marginal adaptive method: How to give nuisance parameters the attention they deserve (no more, no less). J Vis, 13(7), 3. https://doi.org/10.1167/13.7.3

Prins, N., & Kingdom, F. A. A. (2009). Palamedes: Matlab routines for analyzing psychophysical data. http://www.palamedestoolbox.org.

Prins, N., & Kingdom, F. A. A. (2018). Applying the Model-Comparison Approach to Test Specific Research Hypotheses in Psychophysical Research Using the Palamedes Toolbox. Front Psychol, 9, 1250. https://doi.org/10.3389/fpsyg.2018.01250

Rohde, M., van Dam, L. C. J., & Ernst, M. (2016). Statistically Optimal Multisensory Cue Integration: A Practical Tutorial. Multisens Res, 29(4-5), 279–317. https://www.ncbi.nlm.nih.gov/pubmed/29384605

Rosas, P., & Wichmann, F. A. (2011). Cue combination: Beyond optimality. In J. Trommershauser, M. S. Landy, & K.P. Körding (Eds.), Sensory Cue Integration (pp. 144–152). Oxford University Press.

Saunders, J. A., & Chen, Z. (2015). Perceptual biases and cue weighting in perception of 3D slant from texture and stereo information. J Vis, 15(2). https://doi.org/10.1167/15.2.14

Scarfe, P., & Glennerster, A. (2014). Humans use predictive kinematic models to calibrate visual cues to three-dimensional surface slant. Journal of Neuroscience, 34(31), 10394–10401. https://doi.org/10.1523/JNEUROSCI.1000-14.2014

Scarfe, P., & Hibbard, P. B. (2011). Statistically optimal integration of biased sensory estimates. J Vis, 11(7). https://doi.org/10.1167/11.7.12

Schütt, H. H., Harmeling, S., Macke, J. H., & Wichmann, F. A. (2016). Painfree and accurate Bayesian estimation of psychometric functions for (potentially) overdispersed data. Vision Res, 122, 105–123. https://doi.org/10.1016/j.visres.2016.02.002

Serwe, S., Drewing, K., & Trommershauser, J. (2009). Combination of noisy directional visual and proprioceptive information. J Vis, 9(5), 28 21–14. https://doi.org/10.1167/9.5.28

Smeets, J. B., van den Dobbelsteen, J. J., de Grave, D. D., van Beers, R. J., & Brenner, E. (2006). Sensory integration does not lead to sensory calibration [Comparative Study Research Support, Non-U.S. Gov’t]. Proc Natl Acad Sci U S A, 103(49), 18781–18786. https://doi.org/10.1073/pnas.0607687103

Svarverud, E., Gilson, S. J., & Glennerster, A. (2010). Cue combination for 3D location judgements. J Vis, 10(1), 5 1–13. https://doi.org/10.1167/10.1.5

Takahashi, C., Diedrichsen, J., & Watt, S. J. (2009). Integration of vision and haptics during tool use. J Vis, 9(6), 3 1–13. https://doi.org/10.1167/9.6.3

Tassinari, H., & Domini, F. (2008). The intrinsic constraint model for stereo-motion integration. Perception, 37(1), 79–95. https://doi.org/10.1068/p5501

Todd, J. T. (2015). Can a Bayesian analysis account for systematic errors in judgments of 3D shape from texture? A reply to Saunders and Chen. J Vis, 15(9), 22. https://doi.org/10.1167/15.9.22

Todd, J. T., Christensen, J. C., & Guckes, K. M. (2010). Are discrimination thresholds a valid measure of variance for judgments of slant from texture? J Vis, 10(2), 20 21–18. https://doi.org/10.1167/10.2.20

Todd, J. T., & Thaler, L. (2010). The perception of 3D shape from texture based on directional width gradients. J Vis, 10(5), 17. https://doi.org/10.1167/10.5.17

Trommershauser, J., Körding, K. P., & Landy, M. S. (2011). Sensory Cue Integration. Oxford University Press.

Wagner, M. (1985). The metric of visual space. Percept Psychophys, 38(6), 483–495. https://www.ncbi.nlm.nih.gov/pubmed/3834394

Watson, A. B. (2017). QUEST+: A general multidimensional Bayesian adaptive psychometric method. J Vis, 17(3), 10. https://doi.org/10.1167/17.3.10

Watson, A. B., & Pelli, D. G. (1983). QUEST: a Bayesian adaptive psychometric method. Percept Psychophys, 33(2), 113–120. https://doi.org/10.3758/bf03202828

Watt, S. J., Akeley, K., Ernst, M. O., & Banks, M. S. (2005). Focus cues affect perceived depth. J Vis, 5(10), 834–862. https://doi.org/10.1167/5.10.7

Welch, R. B., Bridgeman, B., Anand, S., & Browman, K. E. (1993). Alternating prism exposure causes dual adaptation and generalization to a novel displacement. Percept Psychophys, 54(2), 195–204. https://www.ncbi.nlm.nih.gov/pubmed/8361835

Wichmann, F. A., & Hill, N. J. (2001a). The psychometric function: I. Fitting, sampling, and goodness of fit. Percept Psychophys, 63(8), 1293–1313. https://www.ncbi.nlm.nih.gov/pubmed/11800458

Wichmann, F. A., & Hill, N. J. (2001b). The psychometric function: II. Bootstrap-based confidence intervals and sampling. Percept Psychophys, 63(8), 1314–1329. https://www.ncbi.nlm.nih.gov/pubmed/11800459

Xia, Y., Leib, A. Y., & Whitney, D. (2016). Serial dependence in the perception of attractiveness. J Vis, 16(15), 28. https://doi.org/10.1167/16.15.28

Young, M. J., Landy, M. S., & Maloney, L. T. (1993). A perturbation analysis of depth perception from combinations of texture and motion cues. Vision Res, 33(18), 2685–2696. https://doi.org/Doi10.1016/0042-6989(93)90228-O

Zabulis, X., & Backus, B. T. (2004). Starry night: a texture devoid of depth cues. J Opt Soc Am A Opt Image Sci Vis, 21(11), 2049–2060. https://www.ncbi.nlm.nih.gov/pubmed/15535362

